# Mitochondrial metabolism sustains CD8^+^ T cell migration for an efficient infiltration into solid tumors

**DOI:** 10.1101/2024.01.12.575327

**Authors:** Luca Simula, Mattia Fumagalli, Lene Vimeux, Irena Rajnpreht, Philippe Icard, Dongjie An, Frédéric Pendino, Diane Damotte, Audrey Lupo-Mansuet, Marco Alifano, Marie-Clotilde Alves-Guerra, Emmanuel Donnadieu

## Abstract

The ability of CD8^+^ T cells to infiltrate solid tumors and reach cancer cells is associated with improved patient survival and responses to immunotherapy. Thus, identifying the factors controlling T cell migration in tumors is critical, so that strategies to intervene on these targets can be developed. Although interstitial motility is a highly energy-demanding process, the metabolic requirements of CD8^+^ T cells migrating in a 3D environment remain unclear. Here, we demonstrate that the tricarboxylic acid (TCA) cycle is the main metabolic pathway sustaining human CD8^+^ T cell motility in 3D collagen gels and tumor slices while glycolysis plays a much minor role. Using pharmacological and genetic approaches, we report that CD8^+^ T cell migration depends on the mitochondrial oxidation of glucose and glutamine, but not fatty acids, and both ATP and ROS produced by mitochondria are required for T cells to migrate. Pharmacological interventions to increase mitochondrial activity improve CD8^+^ T cells intra-tumoral migration and CAR T cell recruitment into tumor islets leading to better control of tumor growth in human xenograft models. Our study highlights the rationale of targeting mitochondrial metabolism to enhance the migration and antitumor efficacy of CAR T cells in treating solid tumors.

## INTRODUCTION

Although T cell-based immunotherapies have transformed cancer treatment, only a minority of patients with solid cancers responds to these approaches^1^. Several mechanisms have been proposed to explain the reduced CD8^+^ T cell function within the tumor microenvironment (TME)^2,3^, including a defective CD8^+^ T cell intra-tumoral motility. Indeed, for an effective direct destruction of cancer cells, CD8^+^ T cells must fulfill an active migration allowing them to make contact with malignant cells^4^. In most human solid tumors, tumor cells lie within tumor islets surrounded by a stroma composed of fibers and host-derived suppressive cells^5–7^. Remarkably, accumulating evidence suggests that such TME structure limits CD8^+^ T cells from migrating and contacting their targets. We previously reported that several TME elements, such as pro-tumoral macrophages and extracellular matrix fibers, can sequester CD8^+^ T cells in stromal areas, thus preventing their contact with tumor cells^8–10^. Interestingly, such a hostile TME can also prevent the infiltration of CD8^+^ CAR T cells^11,12^, thus limiting the effectiveness of immunotherapy approaches based on CAR T cell infusion for solid cancer patients. Beyond extracellular determinants, intracellular factors also play an important role in the ability of T cells to migrate and reach tumor cells.

Metabolism sustains T cell migration, which relies on a fine-tuned regulation of cytoskeleton remodeling for which metabolic reactions supply energy in the form of ATP and GTP (for actin and tubulin polymerization). Although ATP is known to support both fast-directional migration^13^ and slow motility^14^ of activated T cells and inhibition of its production reduces T cell motility^15^, very few studies have so far addressed how specific metabolic pathways support T cell migration. Some of them highlighted the role of OXPHOS-produced ATP^13,16^, others that of glycolysis^15,17,18^, whose end-product (lactate) inhibits T cell motility^15^. However, the relative importance of these metabolic pathways in sustaining T cell motility has never been addressed. In addition, most of these studies have been carried out using 2D models (mainly transwell assay or protein-coated surfaces)^13–16^, which are not suited to recapitulate the 3D amoeboid-like motility of CD8^+^ T cells within the TME. This aspect is important since due to nutrients unavailability, the accumulation of waste products, the acid pH and the low oxygen tension, CD8^+^ T cell metabolism can be severely altered within TME^1,19^. Such alterations have already been associated with a defective cytotoxicity, persistence and survival of CD8^+^ T cells, favouring tumor escape^1^. Since motility is a highly energy-demanding process, it is extremely likely that these metabolic alterations could also impair CD8^+^ T cell intra-tumoral motility, thus playing an important role in preventing these cells from reaching and killing tumor cells. However, our lack of knowledge about how metabolic pathways support CD8^+^ T cell 3D motility within the TME prevents us from anticipating how TME metabolic alterations could affect T cell motility and developing effective strategies to overcome these defects, especially for CAR T cell-based strategies.

Therefore, we decided to investigate how metabolism supports CD8^+^ T cell 3D motility using relevant 3D motility models. Our data indicate that (*i*) an effective CD8^+^ T cell 3D motility is supported mainly by glucose- and glutamine-fueled TCA cycle sustaining both ATP and mtROS production from mitochondria and (*ii*) strategies targeting mitochondrial metabolism are effective in increasing the intra-tumoral infiltration of CD8^+^ CAR T cells to help them reach and kill tumor cells in preclinical solid tumor models.

## RESULTS

### Human CD8^+^ T cell 3D motility is mainly supported by the TCA cycle fueled by glucose and glutamine

To investigate the metabolic regulation of CD8^+^ T cell motility in a 3D environment, we evaluated the spontaneous migration of activated CD8^+^ T cells in a 3D collagen gel using time-lapse imaging microscopy. Naïve CD8^+^ T cells were activated (anti-CD3/CD28) and expanded in the presence of IL7+IL15 cytokines (**Fig. 1A**). First, we tested the role of glucose and glutamine, the main energy sources in culture media. During acute deprivation (1h), the absence of glucose, but not glutamine, slightly but significantly reduced 3D motility (**Fig. 1B**). This effect was presumably due to the different concentrations of these nutrients in the culture medium, since increasing glutamine concentration to match glucose one abrogated this difference (**Fig. 1C**). After prolonged deprivation (48h), both nutrients halved 3D motility in a similar way (**Fig. 1D**). These data suggest that glucose and glutamine support similarly 3D motility as they can compensate for each other during a short time, while for longer periods both are similarly required, like it was observed for T cell proliferation (***Fig. S1A***). Interestingly, while only glucose sustains glycolysis, both nutrients may support similarly TCA cycle in T cells, as suggested by CO_2_ flux analysis (***Fig. S1B***). Next, we investigated whether fatty acids (FAs) also sustain 3D motility. Since FAs are almost not present in culture media, we added oleic acid, linoleic acid, or palmitate during the motility assay. Surprisingly, although these FAs have been reported to modulate some functions in T cells^20–24^, we observed no effect on motility (**Fig. 1E**). In line with this, FAO inhibitors etomoxir, perhexiline (Cpt1a transporter) or trimetazidine (3-KAT enzyme) had no impact on motility (**Fig. 1F**). Overall, these data indicate that glucose and glutamine, but not FAs, are required to support CD8^+^ T cell 3D motility under normal conditions.

**Figure 1.**
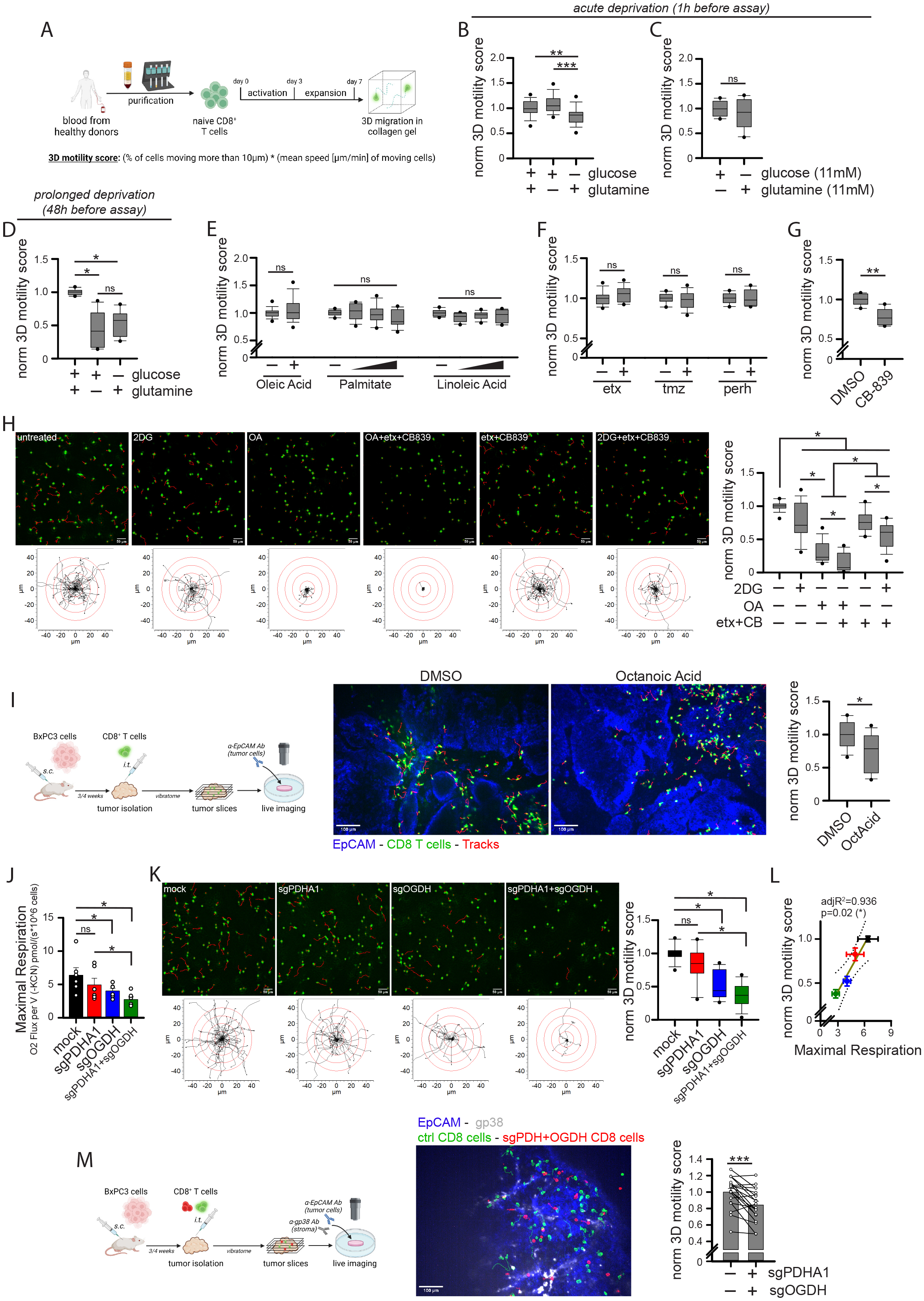
CD8^+^ T cell 3D motility is mainly supported by the TCA cycle fueled by glucose and glutamine but not fatty acids. (**A**) Experimental layout. (**B-H**) Normalized 3D motility Score of activated CD8^+^ T cells in motility medium (see M&M) containing no glucose or no glutamine (B, n=27 movies), either glucose or glutamine at 11mM (C, n=10), starved from glucose or glutamine since 48h before evaluating motility (D, n=12; motility assay also in absence of glucose or glutamine) or in presence of the indicated nutrients or drugs (2DG: 2-deoxy-glucose; etx: etomoxir; tmz: trimetazidine; perh: perhexiline; OA: 6,8-bis(benzylthio)octanoic acid; CB: CB-839) (E, oleic acid n=29, linoleic acid & palmitate n=12; F, etx n=12, tmz & perh n=10; G, n=6; H, n=18). In H, (top) 2D z-stack reconstructions of 3D motility in collagen gel (cells in green, tracks in red) and (bottom) superimposed tracks normalized to their starting coordinates (tracks in black, red circles every 10µm) are shown for each condition. (**I**) Experimental layout on the left. 3D motility of Calcein Green-stained activated CD8^+^ T cells (cells in green, tracks in red) within viable tumor slice (tumor cells in blue, EpCAM) derived from BxPC3/NSG model and incubated with the indicated drugs (n=12 slices). (**J-L**) Measurement of maximal respiration (J, n=6) and normalized 3D motility Score (K, n=15) and correlation between the two parameters (L) in activated CD8^+^ T cells CRISPR/Cas9-edited for the indicated genes. (**M**) Experimental layout on the left. 3D motility of activated Calcein Green-stained control CD8^+^ T cells (green) and Calcein Red-stained CRISPR/Cas9-edited CD8^+^ T cells (sgPDH+sgOGDH) (red) onto viable tumor slice (tumor cells in blue, EpCAM; stroma in gray, gp38) derived from BxPC3/NSG model (n=19). Scale bar, 50µm in H-K and 100µm in I-M.

We next decided to evaluate the impact of pharmacological inhibitors of glycolysis (2DG), TCA cycle (octanoic acid [OA] an inhibitor of pyruvate dehydrogenase [PDH] and oxoglutarate dehydrogenase [OGDH]) and glutaminolysis (CB-839) (see scheme in ***Fig. S1C***). First, we observed that inhibition of glutaminolysis significantly reduced 3D motility (**Fig. 1G**), an effect more pronounced in a medium containing no glucose (***Fig. S1D***). Second, 3D motility was similarly reduced in the presence of 2DG (which inhibits both glycolysis and glucose-derived OXPHOS) or etomoxir + CB-839 combination (whose effect is mainly due to inhibition of glutaminolysis, since etomoxir alone had no effect) (**Fig. 1H** and **Movies S1-6**). Third, a much stronger decrease in 3D motility was observed in the presence of TCA cycle inhibitor OA (**Fig. 1H** and **Movies S1-6**). Fourth, a complete blockade of the TCA cycle by the triple combination of OA + etomoxir + CB-839 (allowing only glycolysis to generate energy) completely blocked 3D motility (less than 10% of initial motility left) (**Fig. 1H** and **Movies S1-6**). Taken together, these data suggest that CD8^+^ T cell 3D motility is mainly supported by the TCA cycle while glycolysis plays a minor role. This is in clear contrast to what is known for T cell proliferation, which relies more on glycolysis-derived energy^25,26^, as we also observed (***Fig. S1E***). Of note, these findings are not limited to cells expanded in the presence of IL-7+IL-15 (which promote mitochondrial metabolism^27^), since similar results were also observed in cells cultured with IL-2 (more glycolysis-prone^28^) (***Fig. S1F-G***). Next, to validate the effects of OA in a more relevant TME-like model, we injected fluorescence-labeled activated CD8^+^ T cells into a fresh tumor fragment obtained from s.c. inoculation of human BxPC3 pancreatic cells into NSG nude mice (termed BxPC3-NSG model). Then, the tumor piece was cut into viable slices and the motility of CD8^+^ T cells was recorded with a spinning disk confocal microscope. Interestingly, slices incubated with OA significantly reduced CD8^+^ T cell 3D motility (**Fig. 1I** and **Movies S7-8**). To confirm these effects, we took advantage of CRISPR/Cas9 approach to edit the OA targets PDH and OGDH. Surprisingly, although efficiently edited (***Fig. S1H***), down-regulation of the PDHA1 gene in T cells (PDHA2 is not expressed) had no impact on either mitochondrial respiration (**Fig. 1J** and ***Fig. S1I***) nor 3D motility (**Fig. 1K** and **Movies S9-12**), indicating the presence of compensatory mechanisms to maintain TCA cycle activity. In contast, OGDH down-regulation (alone or in combination with PDHA1 editing) (***Fig. S1H***) significantly reduces both maximal respiration and 3D motility in CD8^+^ T cells (**Fig. 1J-K** and **Movies S9-12**), an effect also confirmed using OGDH inhibitor succinyl-phosphonate (***Fig. S1J***). Remarkably, under these conditions, we observed an almost perfect correlation between maximal mitochondrial respiration and 3D motility in collagen gel (**Fig. 1L**), further corroborating the notion that mitochondrial metabolism is the main player supporting 3D motility. Furthermore, CRISPR/Cas9-mediated double PDHA1/OGDH knock-down significantly reduced CD8^+^ T cell intra-tumoral motility as assessed using tumor slice derived from the BXPC3-NSG model (**Fig. 1M** and **Movie S13**). Last, we observed that CD8^+^ T cell 3D motility in collagen gel can be increased by forcing glucose-derived pyruvate to enter TCA cycle through administration of (*i*) lactate dehydrogenase (LDHi) and PDH kinase (PS10) inhibitors or (*ii*) a PDH activator (α-lipoic acid) (***Fig. S1K-L***).

Overall, these data indicate that CD8^+^ T cell 3D motility is mainly supported by mitochondrial metabolism, and more specifically by the TCA cycle fueled by glucose and glutamine.

### Glucose fuels human CD8^+^ T cell 3D motility mainly via the TCA cycle and to a lesser extent via glycolysis

Our data (**Fig. 1**) suggested that glucose sustains CD8^+^ T cell motility mainly via OXPHOS and poorly via glycolysis. To further clarify the relative importance of these two pathways in supporting CD8^+^ T cells 3D motility, we performed the following experiments.

First, to force cells to rely on OXPHOS-derived ATP, we cultured CD8^+^ T cells in the presence of glucose or galactose for 3 days. Of note, galactose cannot provide ATP via glycolysis but only via OXPHOS^29,30^. While 3D motility was minimally impacted by the replacement of glucose by galactose (**Fig. 2A** and **Movies S14-15**), proliferation was strongly impaired (**Fig. 2B**), consistent with the notion that T cell proliferation, but not motility, requires glycolytic ATP.

**Figure 2.**
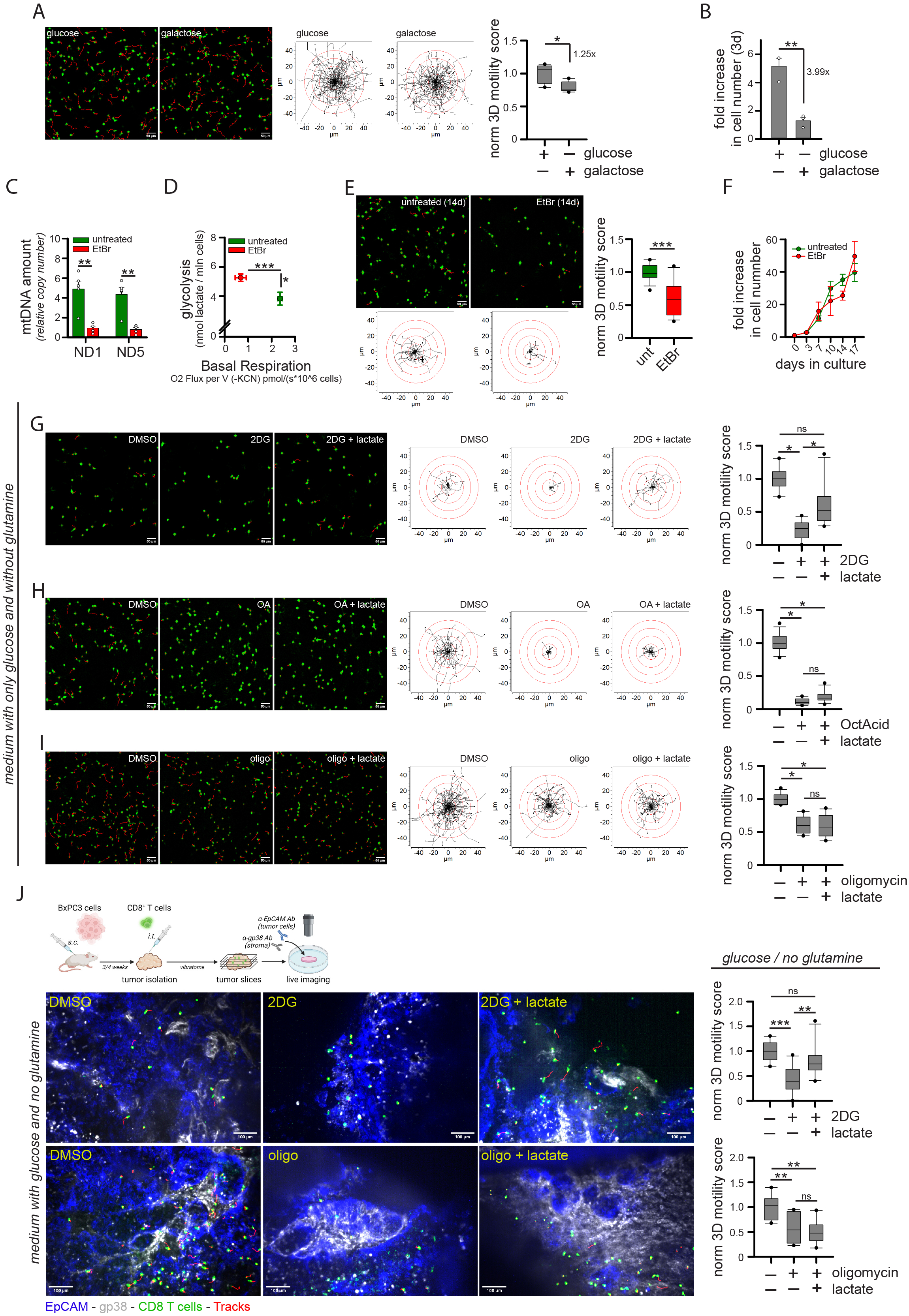
Glucose fuels CD8^+^ T cell 3D motility mainly through the TCA cycle and poorly through glycolysis. (**A-B**) Normalized 3D motility Score (A, n=5) and fold increase in cell number (B, n=3) of activated CD8^+^ T cells cultured since 72h before motility assay in medium containing 11mM glucose or galactose (n=5). (**C-F**) Activated CD8^+^ T cells were cultured for at least 14d in control medium (untreated) or in the presence of low-dose ethidium bromide (EtBr). After 14d, the following parameters were measured: mtDNA amount (C, 2^-ΔCt^ values between mtDNA genes ND1 or ND5 and nuclear gene 18S; n=5), energy plot (D, raw measurements are reported in Figure S2A-B), normalized 3D motility Score (E, n=15), fold increase in cell number (F, n=3). (**G-I**) Normalized 3D motility Score of activated CD8^+^ T cells in the presence of 2-deoxy-glucose (2DG; G, n=10), 6,8-bis(benzylthio)octanoic acid (OA/OctAcid; n=12) and oligomycin (oligo; I, n=12) with or without lactate in a medium containing only glucose and no glutamine. (**J**) Experimental layout on the top left. 3D motility of Calcein Green-stained activated CD8^+^ T cells (cells in green, tracks in red) within viable tumor slices (tumor cells in blue, EpCAM; stroma in gray, gp38) derived from BxPC3/NSG model and incubated with the indicated drugs or nutrients (2DG, n=10; oligo, n=8). Scale bar, 50µm in A-E-G-H-I and 100µm in J.

Second, we cultured CD8^+^ T cells in the presence of a low dose of ethidium bromide (EtBr) to deplete mtDNA^31^, forcing the cells to rely on glycolytic ATP. 2-week EtBr treatment significantly reduced mtDNA amount (**Fig. 2C**) and mitochondrial respiration while increasing glycolysis (**Fig. 2D**, see also ***Fig. S2A-B***) and mtROS levels (***Fig. S2C***). In contrast to what was observed with galactose, EtBr-treated CD8^+^ T cells showed a strong reduction in 3D motility (**Fig. 2E** and **Movies S16-17**) but no effect on cell proliferation (**Fig. 2F**).

Third, we evaluated the ability of several nutrients to rescue motility when glycolysis or OXPHOS were inhibited in a medium containing only glucose and no glutamine. Under this condition, the addition of 2DG strongly inhibited 3D motility (**Fig. 2G** and **Movies S18-20**). However, 3D motility was restored by the addition of lactate, which bypasses upstream glucose metabolism and directly fuels the mitochondrial TCA cycle (**Fig. 2G** and **Movies S18-20**). In contrast, lactate failed to rescue the inhibition of 3D motility due to TCA-cycle inhibitor OA (**Fig. 2H** and **Movies S21-23**) or OXPHOS inhibitor oligomycin (**Fig. 2I** and **Movies S24-26**). A similar rescue of 2DG-but not oligomycin-mediated inhibition was observed using additional nutrients downstream of glycolysis (pyruvate and acetate) and cell-permeable derivatives of TCA cycle intermediates (dimethyl-succinate and dimethyl-α-ketoglutarate) (***Fig S2D-E***). Of note, the ability of lactate to rescue 2DG-but not oligomycin-induced reduction in motility was also observed in tumor slices derived from the BxPC3-NSG model (**Fig. 2J** and **Movies S27-32**). Last, the administration of nutrients such as lactate, acetate, or pyruvate increased 3D motility even in complete medium containing both glucose and glutamine (***Fig. S2F***), presumably by providing an extra boost to TCA cycle and OXPHOS machinery.

In sum, these data indicate that (*i*) glucose sustains CD8^+^ T cell 3D intra-tumoral motility mainly via OXPHOS and not glycolysis and (*ii*) several nutrients (including lactate) could be used by CD8^+^ T cells to support their motility in the absence of glucose.

### Human CD8^+^ T cell mitochondrial metabolism and 3D motility are correlated in different contexts

Given the important role of mitochondrial metabolism in supporting CD8^+^ T cell 3D motility, we wondered whether these two parameters correlate in CD8^+^ T cells under different conditions.

First, we decided to compare CD8^+^ T cells expanded in the presence of IL-2 or IL-7+IL-15 (IL-7/15), since these cytokines differentially affect the glycolysis/OXPHOS rate in T cells^27,28^. IL-7/15 CD8^+^ T cells showed a similar glycolytic rate but increased mitochondrial respiration (**Fig. 3A** and ***Fig. S3A-B***) and higher mitochondrial mass and glucose uptake with similar levels of mitochondrial membrane potential (MMP) (***Fig. S3C***) compared to IL-2 cells. In line with our data, such increased mitochondrial activity correlated with a superior 3D motility in collagen gel (**Fig. 3B** and **Movies S33-34**). Under these conditions, we also found that 3D motility correlated strongly with several OXPHOS parameters, such as mitochondrial mass, mitochondrial membrane potential (MMP) and expression of the TCA-cycle enzyme isocitrate-dehydrogenase-2 (IDH2), as well as with global glucose uptake and expression of glucose receptor GLUT1 (***Fig. S3D***). On the contrary, 3D motility was not or negatively correlated with the level of expression of glycolytic enzymes phospho-fructokinase-2 (PFK2) and hexokinase-1 (HK-1), as well as with the monocarboxylate transporter-1 (MCT-1) and the pentose-phosphate pathway (PPP) regulator glucose-6-phosphate-dehydrogenase (G6PD) (***Fig. S3D***). The higher motility of IL7+15 CD8^+^ T cells compared to IL2 cells was also confirmed in tumor slices derived from the BxPC3-NSG model (**Fig. 3C** and **Movie S35**). This enhancement was mainly due to IL-15 (***Fig. S3E-F***), in line with its prominent role in supporting mitochondrial metabolism^27^.

**Figure 3.**
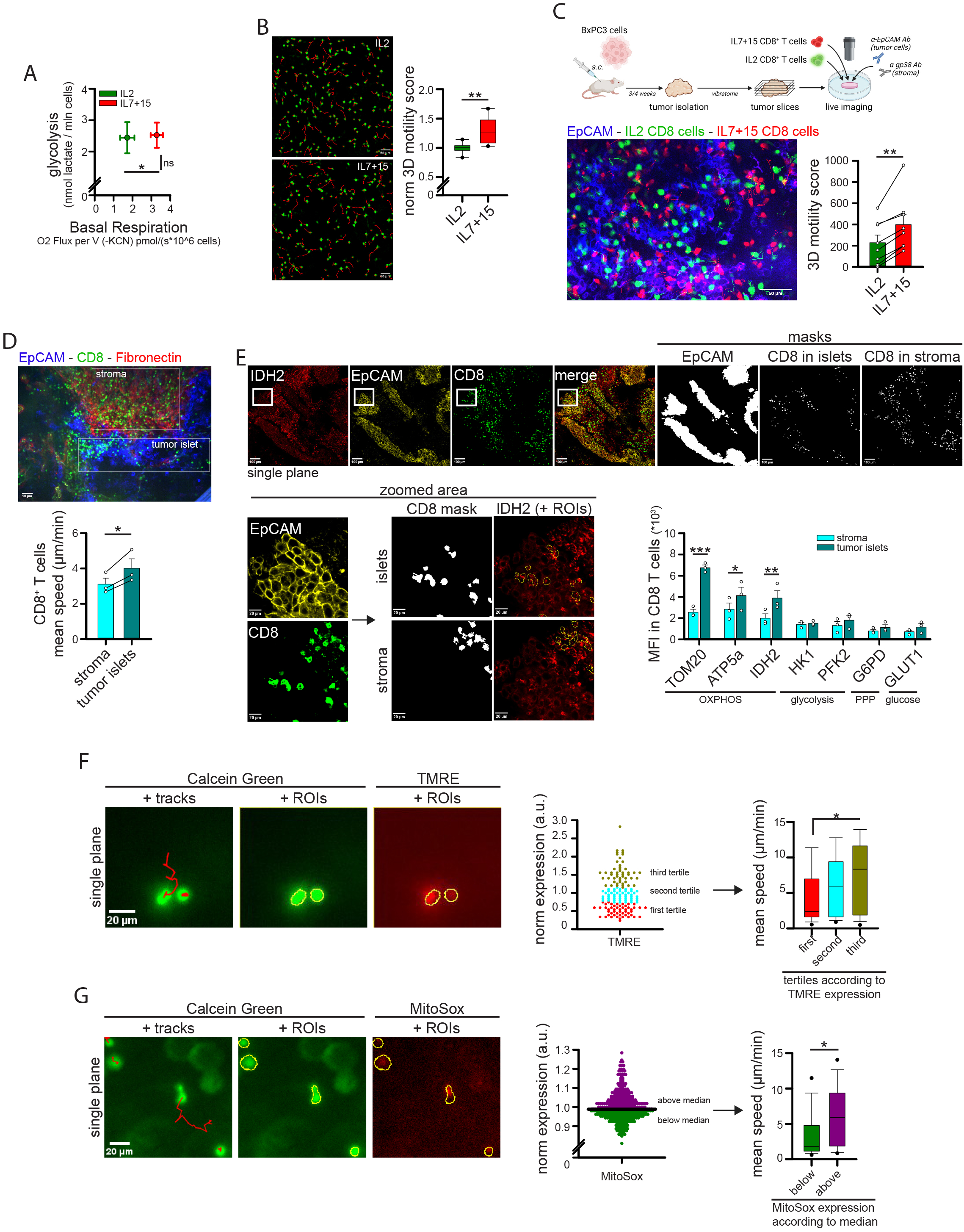
CD8^+^ T cell mitochondrial metabolism and 3D motility correlate across multiple contexts. (**A-B**) CD8^+^ T cells have been activated (3d) and expanded (4d) in the presence of IL-2 or IL-7+IL-15 (IL7+15). After 7 days, the energy plot (A, raw measurements are reported in Figure S3A-B), and normalized 3D motility Score (B, n=8) were calculated. (**C**) Experimental layout on top. 3D motility of activated Calcein Green-stained IL2 CD8^+^ T cells (green) and Calcein Red-stained IL7+15 CD8^+^ T cells (red) overlaid onto viable tumor slices (tumor cells in blue, EpCAM) derived from BxPC3/NSG model (n=8). (**D**) 3D motility of resident CD8^+^ T cells (anti-CD8, green) in viable tumor slices (tumor cells in blue, EpCAM; stroma in red, fibronectin) derived from human NSCLC biopsies (n=3). (**E**) Mean level of expression (MFI) of the indicated proteins calculated in CD8^+^ T cells located in tumor islets or stroma of NSCLC biopsies, as identified from immunofluorescent stainings (see methods for details) from 3 different donors (confocal images on top and bottom left). The panel shows representative images of IDH2 staining only. (**F-G**) Activated CD8^+^ T cells have been stained with Calcein Green (green) and TMRE (F, red) or MitoSox (G, red) and 3D motility was assessed in collagen gel. For each cell, motility was evaluated by TrackMate analysis (tracks in red) and TMRE or MitoSox MFI was calculated at the first timepoint (cells were identified using ImageJ ROIs). Cells were stratified according to TMRE or MitoSox expression and motility was calculated in each subgroup (graphs on the right) (F, n=150 cells; G, n=420 cells). Scale bar, 50µm in B-C-D, 20-100µm in E and 20µm in F-G.

Second, to compare T cell motility in the absence of culturing cytokines, we evaluated 3D motility and the level of several metabolic parameters in freshly isolated human peripheral blood T (hPBT) cells stained with anti-CD8 and anti-CD45RA Abs to distinguish 4 populations (CD8 or CD4 either naïve-like 45RA^pos^ or memory-like 45RA^neg^). Once again, we observed that the subpopulation migrating more efficiently (*i.e.*, CD8^neg^45RA^neg^ cells, CD4 memory-like) exhibited higher mitochondrial mass and MMP, while no selective increase in glycolysis-associated parameters (***Fig. S3G***).

Third, we previously reported that CD8^+^ T cells localizing in tumor islets of non-small cell cancer (NSCLC) biopsies move faster than CD8^+^ T cells in the surrounding stroma^10^, an observation which we confirmed (**Fig. 3D**). We therefore wondered whether this higher motility correlates with an increase in mitochondria-associated parameters. We observed that CD8^+^ T cells located within tumor islets showed higher expression of several mitochondrial proteins (TOM20, IDH2 and ATP5a) compared to CD8^+^ T cells located in the surrounding stroma (**Fig. 3E**). In contrast, no differences were observed for key glycolytic enzymes, such as HK1 and PFK2 (**Fig. 3E**). These data indicate that 3D motility and mitochondrial parameters correlate when comparing CD8^+^ T cell subpopulations infiltrating NSCLC tissue.

Last, we asked whether the correlation between motility and mitochondrial metabolism holds true also when comparing different cells within the same CD8^+^ population, *i.e.*, if the cells moving faster also show higher mitochondrial activity. To answer this point, we evaluated 3D motility of CD8^+^ T cells stained with TMRE or MitoSox (to monitor MMP and mtROS respectively, two proxies of mitochondrial activity in cells). As shown in **Fig. 3F-G**, CD8^+^ T cells migrating faster also showed higher levels of MMP and mtROS amount compared to slower cells.

Overall, our observations indicate that mitochondrial metabolism and 3D motility are positively correlated in CD8^+^ T cells in multiple contexts.

### TCA cycle sustains ATP and ROS production from mitochondria to support human CD8^+^ T cell 3D motility

The highly energetic process of CD8^+^ T cell migration in 2D^13,14^ and 3D environments helps explain the high requirement for mitochondria, which are the main sources of cellular ATP. Indeed, CD8^+^ T 3D motility is progressively more inhibited after 1h of incubation with increasing doses of oligomycin (**Fig. 4A**). However, we noticed a strong reduction in 3D motility between 1µM and 10µM doses, which cannot be explained by differences in ATP production, since OXPHOS was already efficiently blocked by 1µM of oligomycin (**Fig. 4B**). Both doses showed no toxic effects on mitochondrial functioning, since mitochondrial respiration correctly re-increased after CCCP administration, indicating unaltered ETC functionality (***Fig. S4A*** and **Fig. 4B**). Interestingly, we noticed an abrupt decrease in mtROS amount between these two doses of oligomycin, correlating with the reduction in 3D motility (**Fig. 4C**) and suggesting that also mtROS might positively sustain 3D motility. ROS have pleiotropic roles in the regulation of cell migration^32–35^, particularly the amoeboid-like motility typical of T cells within 3D environment^36,37^. To better clarify the role of ATP and mtROS in CD8^+^ T cell 3D migration, we performed additional experiments.

**Figure 4.**
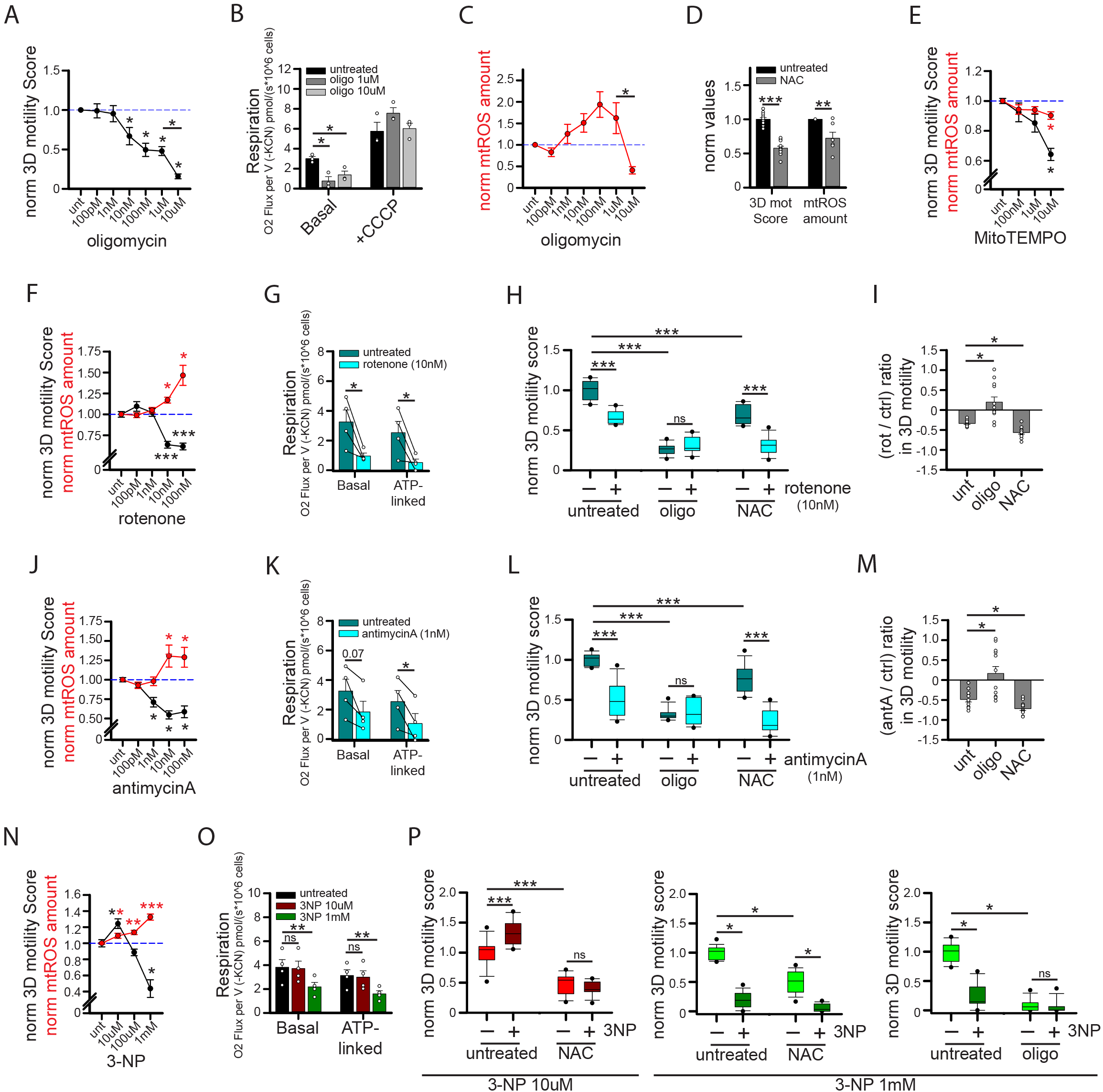
Mitochondria sustain CD8^+^ T cell 3D motility by providing both ATP and mtROS. (**A-C**) Normalized 3D motility Score (A, n=8), basal and CCCP-induced respiration (B, n=3) and normalized mtROS amount (MitoSox/FSC ratio; C, n=7) of activated CD8^+^ T cells in the presence of the indicated doses of oligomycin. (**D-E**) Normalized 3D motility Score (D, n=12; E, n=15) and mtROS amount (MitoSox/FSC ratio; D, n=6; E, n=6) of activated CD8^+^ T cells in the presence of N-acetylcysteine (NAC) (in D) or MitoTEMPO (in E). (**F-M**) Normalized 3D motility Score (F, n=16; J, n=16; N, n=12), mtROS amount (MitoSox/FSC ratio; F, n=7; J, n=8; N, n=6) and basal and ATP-linked respiration (G, n=4; K, n=4; O, n=4) of activated CD8^+^ T cells in the presence of the indicated doses of rotenone (F-K), antimycinA (J-M) or 3-nitropropionic acid (3-NP) (N-P). In H, L and P, motility was evaluated also in the presence of N-acetylcysteine (NAC) or oligomycin (oligo) (H-L, n=12; P, 3-NP 10µM, n=12; oligo + 3-NP 1mM, n=12; NAC + 3-NP 1mM, n=18) and the relative (rotenone/control) or (antimycinA/control) relative changes in 3D motility Score for each condition were reported in I, M. Please note that the same controls were used for data in Figures 4G and 4K since performed in the same experiments (data were split to improve clarity of the presentation).

First, to modulate ATP levels without affecting mtROS, we incubated migrating cells with the uncoupling agent CCCP for 1h. This led to a reduction in MMP (***Fig. S4B***) and ATP-linked mitochondrial respiration (***Fig. S4C***) without affecting mtROS level (***Fig. S4D***). Under this condition, CD8^+^ T cell 3D motility was strongly inhibited (***Fig. S4E***). To further confirm the correlation between ATP and 3D motility, we expressed the ATP biosensor PercevalHR^38,39^ in CD8^+^ T cells and analyzed their motility in a collagen gel. Motile cells showed higher levels of PercevalHR signal (a proxy for ATP amount) compared with immotile cells (***Fig. S4F***). These data confirm the role of mitochondrial ATP in supporting CD8^+^ T 3D motility.

Second, to investigate mtROS, we first tested the effect of ROS scavengers on 3D motility. Of note, both pan-ROS scavenger N-acetylcysteine (NAC) and mtROS-specific scavenger MitoTEMPO significantly reduced 3D motility at doses affecting mtROS amount (**Fig. 4D-E**), suggesting that mtROS may have a positive role in supporting CD8^+^ T cell 3D motility. We then wondered whether increasing mtROS amount could lead to an increase in 3D motility. For this purpose, we incubated CD8^+^ T cells with rotenone, an inhibitor of the ETC-I complex, during 3D migration in collagen gel. Surprisingly, at a dose capable of increasing mtROS amount (10nM), rotenone inhibited CD8^+^ T cell 3D migration (**Fig. 4F**). However, we noticed that at this dose rotenone also inhibited basal and ATP-linked respiration (**Fig. 4G**), and thus ATP production, which is known to positively support CD8^+^ T cell 3D migration. Interestingly, the rotenone-induced reduction in cell migration was abrogated in presence of oligomycin whereas it was still observed (even at stronger level) with the ROS scavenger NAC (**Fig. 4H-I**). These data indicate that the rotenone-induced inhibition of 3D motility was mainly due to the ATP depletion and not to increased mtROS amount. Also, the stronger effect of rotenone-induced reduction in 3D migration in the presence of NAC (**Fig. 4I**) suggests that mtROS may play a positive role counteracting the reduction in motility due to ATP depletion in this condition. We observed similar results with antimycin-A, an inhibitor of the ETC-III complex (**Fig. 4J-M**). In this case, the key role of ATP depletion was even more evident, since antimycin-A began to inhibit motility at 1nM dose, while significantly affecting mtROS amount (at least when assessed by MitoSox staining) only at higher doses (**Fig. 4J**). Last, we tested the ETC-II complex inhibitor 3-nitropropionic acid (3-NP). Noteworthy, 3-NP increased both mtROS amount and 3D motility (**Fig. 4N**) without affecting mitochondrial respiration and ATP production (**Fig. 4O**) at 10µM, whereas at higher doses (1mM) mtROS amounts are further increased but motility and ATP production are decreased (**Fig. 4N-O**). Interestingly, the increase in cell motility due to 3-NP at 10µM was abrogated by NAC treatment (indicating its dependence on mtROS), while the reduction observed at 1mM was still observed in presence of NAC (when ROS are scavenged) and only disappeared upon oligomycin treatment (indicating its dependence on ATP depletion) (**Fig 4P**). We also confirmed these results using another ETC-II complex inhibitor, namely atpenin-A5 (Atp-A5). Also in this case, a low dose of Atp-A5 increased CD8^+^ T cell 3D migration in a NAC-dependent way, while at higher doses Atp-A5 reduced motility because of ATP depletion despite the concomitant increase in mtROS amount (***Fig. S4G-H***).

Taken together, our data suggest that both mitochondrial ATP and mtROS support CD8^+^ T cell 3D motility. However, ATP levels play a prominent role, and mtROS can only further sustain motility if their accumulation does not affect concomitantly the OXPHOS-dependent ATP production.

### Mitochondrial metabolism also sustains CD4^+^ T cell 3D motility

We wondered whether mitochondrial metabolism also plays a role in supporting CD4^+^ T cell 3D motility like what has been observed for CD8^+^ cells. To answer this question, naïve CD4^+^ T cells were activated and cultured like CD8^+^ T cells and their motility assessed in a 3D collagen gel. We found that: (*i*) prolonged glucose or glutamine impacted similarly on 3D motility (***Fig. S5A***); (*ii*) 2DG and CB-839 inhibitors reduced similarly 3D motility, while etomoxir had no effect (***Fig. S5B***); (*iii*) the strongest inhibition of 3D motility was observed in presence of OA (***Fig. S5B***); (*iv*) complete blockade of TCA cycle completely abrogated 3D motility (***Fig. S5B***); and (*v*) both ATP and mtROS production sustained 3D motility (***Fig. S5C-D***).

Overall, these data suggest that glucose- and glutamine-fueled TCA cycle generating OXPHOS-derived ATP and mtROS is the main metabolic pathway sustaining 3D motility also in CD4^+^ T cells.

### Pharmacological approaches promoting mitochondrial metabolism in human CD8^+^ T cells increase 3D intra-tumoral motility

We then developed strategies promoting mitochondrial metabolism to enhance CD8^+^ T cell 3D migratory ability. To this aim, we first activated and cultured CD8^+^ T cells in the presence of rapamycin (Rapa-CD8 cells), an mTOR inhibitor known to sustain mitochondrial metabolism in immune cells^40,41^ (see scheme in **Fig. 5A**). 1-week treatment with rapamycin did not affect viability (***Fig. S6A***) but significantly increased mitochondrial mass, as assessed by quantification of mtDNA amount (**Fig. 5B**), mitochondrial mass, MMP (***Fig. S6B***) and levels of ETC complexes (***Fig. S6C***). Consequently, Rapa-CD8 cells showed a higher maximal respiratory capacity (**Fig. 5C**), while reducing glycolysis (***Fig. S6D***). Interestingly, these cells also showed an improved 3D motility in collagen gel (**Fig. 5D** and **Movies S36-37**), an effect dependent on the increased mitochondrial activity, since oligomycin abrogated it (**Fig. 5E** and **Movies S38-41**). Remarkably, in contrast to what was observed for ctrl-CD8 cells, etomoxir significantly reduced the 3D motility of Rapa-CD8 cells (**Fig. 5F**), suggesting that rapamycin also opened FAs usage to sustain motility. Rapa-CD8 cells also showed better intra-tumoral motility compared with ctrl-CD8 cells, as observed using tumor slices derived from the BxPC3-NSG model (**Fig. 5G** and **Movie S42**). Moreover, to monitor 3D lymphocyte infiltration into a solid environment, CD8^+^ T cells were added adjacent to a BxPC3 tumor cells-filled collagen gel and their ability to infiltrate evaluated. Remarkably, Rapa-CD8 cells infiltrated the 3D environment faster than ctrl-CD8 cells (**Fig. 5H**). Consistent with previous studies^42,43^, rapamycin treatment also increased CD8^+^ T cell differentiation towards a more primitive (Tscm-like) state (***Fig. S6E***) and reduced IFNγ and perforin production (but not granzyme B and TNFα) upon restimulation (***Fig. S6F***), presumably due to inhibition of glycolysis^44^.

**Figure 5.**
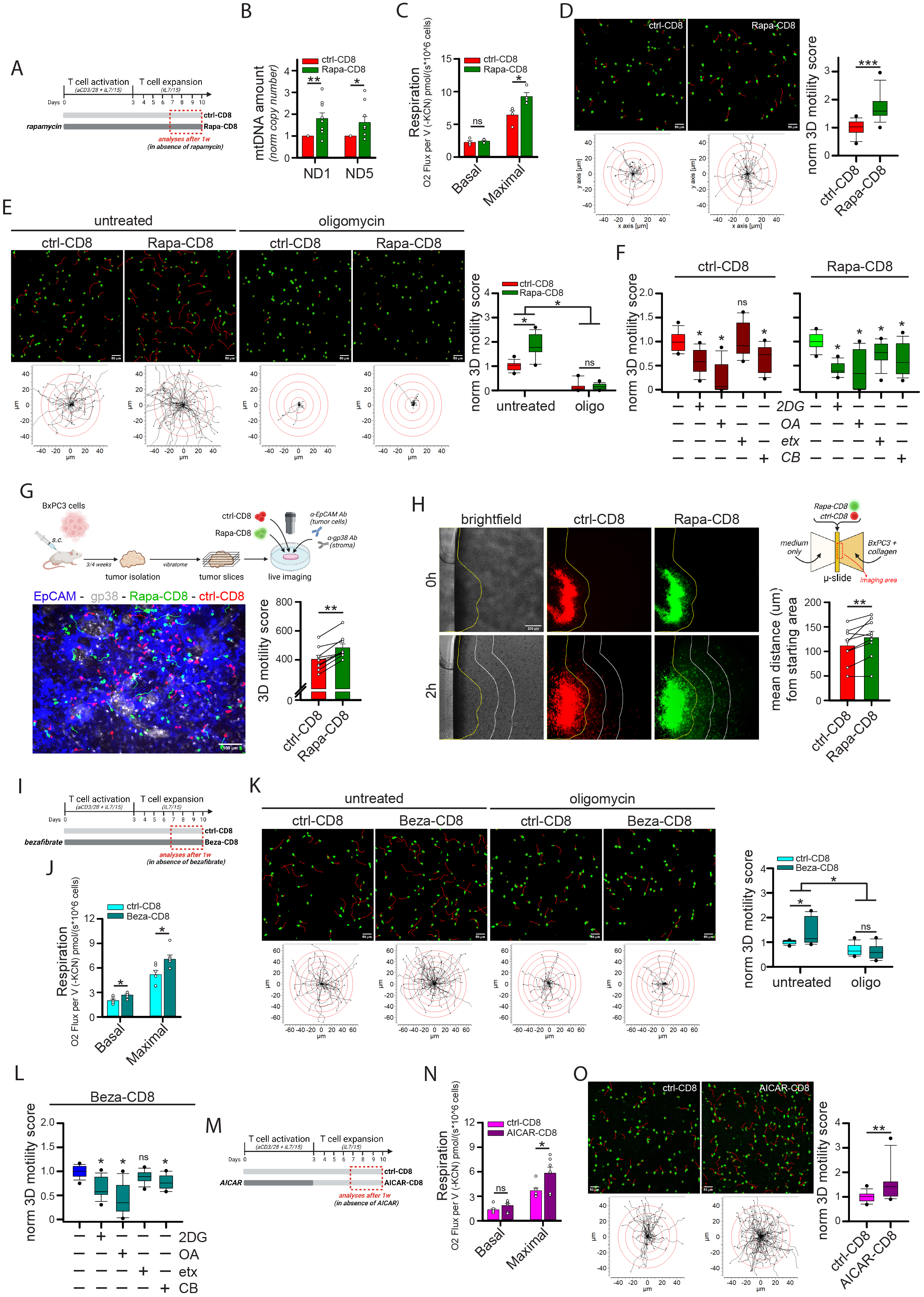
Drugs promoting mitochondrial metabolism increase CD8^+^ T cell 3D motility. (**A-F**) Experimental layout in A. CD8^+^ T cells have been cultured in ctrl condition (ctrl-CD8) or in the presence of 10nM rapamycin (Rapa-CD8). After 1w, rapamycin was removed on the day of analysis and the following parameters were measured: mtDNA amount (normalized 2^-ΔCt^ values between mtDNA genes ND1 or ND5 and nuclear gene 18S; B, n=9), basal and maximal respiration (C, n=4), normalized 3D motility Score in the presence or not of oligomycin (D, n=30; E, n=15). In F (ctrl-CD8, n=12; Rapa-CD8, n=15), the relative 3D motility Score is reported after normalizing to 1 the mean score for both ctrl-CD8 and Rapa-CD8 cells (2DG: 2-deoxy-glucose, bOA: 6,8-bis(benzylthio)octanoic acid; etx: etomoxir; CB: CB-839). (**G**) Experimental layout on the left. 3D motility of activated Calcein Green-stained ctrl-CD8 cells (shown in red for consistency) and Calcein Red-stained Rapa-CD8 cells (shown in green for consistency) overlaid onto viable tumor slices derived from BxPC3/NSG model (tumor cells in blue, EpCAM; stroma in gray, gp38) (n=9). (**H**) Infiltration of Calcein Red-stained ctrl-CD8 cells (red) and Calcein Green-stained Rapa-CD8 cells (green) into a collagen solution containing human BxPC3 tumor cells (scheme on the top right, see methods for details). Yellow lines identify starting position of cells at 0h. White lines represent 200µm increments. The mean distance from starting area for the two populations is reported in the graph (n=9 slides). (**I-L**) Experimental layout in I. CD8^+^ T cells have been cultured in ctrl condition (ctrl-CD8) or in the presence of 10µM bezafibrate (Beza-CD8). After 1w, bezafibrate was removed and the following parameters were measured: basal and maximal respiration (J, n=6) and normalized 3D motility Score in the presence or not of oligomycin (K, n=12). In L (n=15), the relative 3D motility Score is reported after normalizing to 1 the mean score for Beza-CD8 cells (as in 5F). (**M-O**) Experimental layout in M. Activated CD8^+^ T cells have been cultured in ctrl condition (ctrl-CD8) or in the presence of 1mM AICAR (AICAR-CD8). After 3d, AICAR was removed, and the cells expanded in the control condition. The following parameters were measured starting from 7d: basal and maximal respiration (N, n=7) and normalized 3D motility Score (O, n=15). Scale bar, 50µm in D-E-K-O, 100µm in G and 200µm in H.

*In vitro* treatment with rapamycin could therefore be an effective strategy for enhancing the infiltration and intra-tumoral motility of CD8^+^ T cells. To ensure that such an effect could be still maintained several days after rapamycin withdrawal, CD8^+^ T cells treated *in vitro* for 10 days were deprived of rapamycin and their functions evaluated some days after (see scheme in ***Fig. S6G***). Of note, 4/7 days after rapamycin withdrawal, cytokine production was not only restored but even increased in Rapa-CD8 cells (***Fig. S6H***), and these cells retained a better 3D motility compared with ctrl-CD8 cells (***Fig. S6I***). Moreover, although rapamycin treatment reduced *in vitro* CD8^+^ T cell expansion (***Fig. S6J***), this effect was mainly limited to the first few days of culture and disappeared after rapamycin removal (***Fig. S6K***), suggesting that expansion of Rapa-CD8 cells in the host may be similar to control cells. Last, Rapa-CD8 cells also showed an enhanced 3D motility in a different type of collagen (telo-collagen) compared to ctrl-CD8 cells (***Fig. S6L***).

Next, we activated and cultured CD8^+^ T cells in the presence of bezafibrate (Beza-CD8 cells), which can support mitochondrial metabolism in T cells^45^ (see scheme in **Fig. 5I**). Bezafibrate treatment did not affect T cell viability (***Fig. S7A***) but increased mitochondrial respiration (**Fig. 5J**), although this effect was not associated to an increase in mitochondrial mass (***Fig. S7B***). Of note, Beza-CD8 cells showed an oligomycin-dependent increase in 3D motility (**Fig. 5K** and **Movies S43-46**). Unlike Rapa-CD8 cells, Beza-CD8 cells did not acquire the ability to use FAs to fuel their motility (**Fig. 5L**), and their glycolysis and *in vitro* proliferation were not affected (***Fig. S7C-D***). The enhanced 3D motility was also observed using telo-collagen gel (***Fig. S7E***).

Third, we activated CD8^+^ T cells in the presence of AICAR, an activator of the energy regulator AMPK^46^ (AICAR-CD8 cells), for 3 days before expanding them *in vitro* in the absence of the drug (a more prolonged treatment was toxic) (see scheme in **Fig. 5M**). Like bezafibrate, AICAR had no impact on cell viability (***Fig. S7F***), mitochondrial mass (***Fig. S7G***) or glycolysis (***Fig. S7H***) but significantly increased maximal respiration (**Fig. 5N**) and 3D motility (**Fig. 5O** and **Movies S47-48**) in CD8^+^ T cells. Proliferation was reduced similarly to what was observed for rapamycin (***Fig. S7I***).

Last, βNAD treatment (***Fig. S7J***), which modulates mitochondrial metabolism in T cells^47^, increased ATP-linked respiration (***Fig. S7K***) and 3D motility (***Fig. S7L***) in CD8^+^ T cells. Remarkably, the effects of rapamycin, AICAR and βNAD were due to a metabolic reprogramming since administration of these drugs during the motility assay had no affect or even inhibited 3D motility (***Fig. S7M***).

Overall, these data indicate that CD8^+^ T cell 3D motility and intra-tumoral infiltration can be enhanced using pharmacological strategies targeting mitochondrial metabolism.

### Rapamycin-treated human CD8^+^ CAR T cells exhibit superior mitochondrial metabolism and 3D intra-tumoral motility

The ability to equip CD8^+^ CAR T cells with better infiltration and intra-tumoral motility could significantly improve the efficacy of CAR-based immunotherapies against solid cancers. Therefore, we first decided to investigate if mitochondrial metabolism is also the key driver of 3D motility in CD8^+^ CAR T cells. To this aim, CD8^+^ T cells were transduced with lentiviral particles to generate anti-EGFR CD8^+^ CAR T cells and their 3D motility was assessed in collagen gels. Consistent with what was observed in normal cells, glucose and glutamine similarly supported CD8^+^ CAR T cells 3D motility (***Fig. S8A***) and TCA-cycle inhibitor OA strongly reduced it (***Fig. S8B***). Also, 2DG and CB-839 had a similar effect, while etomoxir had no impact (***Fig. S8B***). Motility was also strongly suppressed by oligomycin (***Fig. S8C***). Finally, CRISPR/Cas9-mediated editing of OA targets PDHA1 and OGDH (***Fig. S8D***) led to a significant decrease in 3D motility (***Fig. S8E***). Overall, these data indicate that mitochondrial metabolism strongly supports also CD8^+^ CAR T cell 3D motility.

Next, we tested whether pharmacological strategies enhancing the mitochondrial activity of CD8^+^ CAR T cells could also increase intra-tumoral infiltration. Among the different drugs tested, we selected rapamycin since this drug produced the strongest effects in terms of motility and increase in mitochondrial mass. CD8^+^ T cells were activated in presence of rapamycin, transduced with anti-EGFR CAR lentiviral particles (Rapa-CD8^CAR^ cells), and further expanded for up to 10 days in the presence of the drug, which was removed on the day of the analysis (see scheme in **Fig. 6A**). Rapamycin treatment had no impact on CAR expression (***Fig. S9A***) but significantly increased maximal respiration (**Fig. 6B**), while reducing glycolysis (***Fig. S9B***). Remarkably, Rapa-CD8^CAR^ cells showed enhanced 3D motility in collagen gel (**Fig. 6C** and **Movies S49-50**), an effect mainly due to a higher mean speed of moving cells (***Fig. S9C***). To test the motility of these cells in a solid-TME model, NSG mice were i.v. inoculated with human A549 lung tumor cells (see scheme in **Fig. 6D**), which rapidly populated the lungs generating a relevant TME structure composed of medium-size tumor islets surrounded by a large stroma. After 3 weeks, lungs were excised and cut into viable slices. Interestingly, Rapa-CD8^CAR^ cells showed improved intra-tumoral motility when overlaid onto the lung tumor slices compared to ctrl-CD8^CAR^ cells (**Fig. 6D** and **Movies S51-52**). We also evaluated the ability of CD8^+^ CAR T cells to migrate across a TNFα-activated HUVEC monolayer in a transwell assay as a model of T cell extravasation. Rapa-CD8^CAR^ cells migrated through the HUVEC monolayer faster than control cells (**Fig. 6E**).

**Figure 6.**
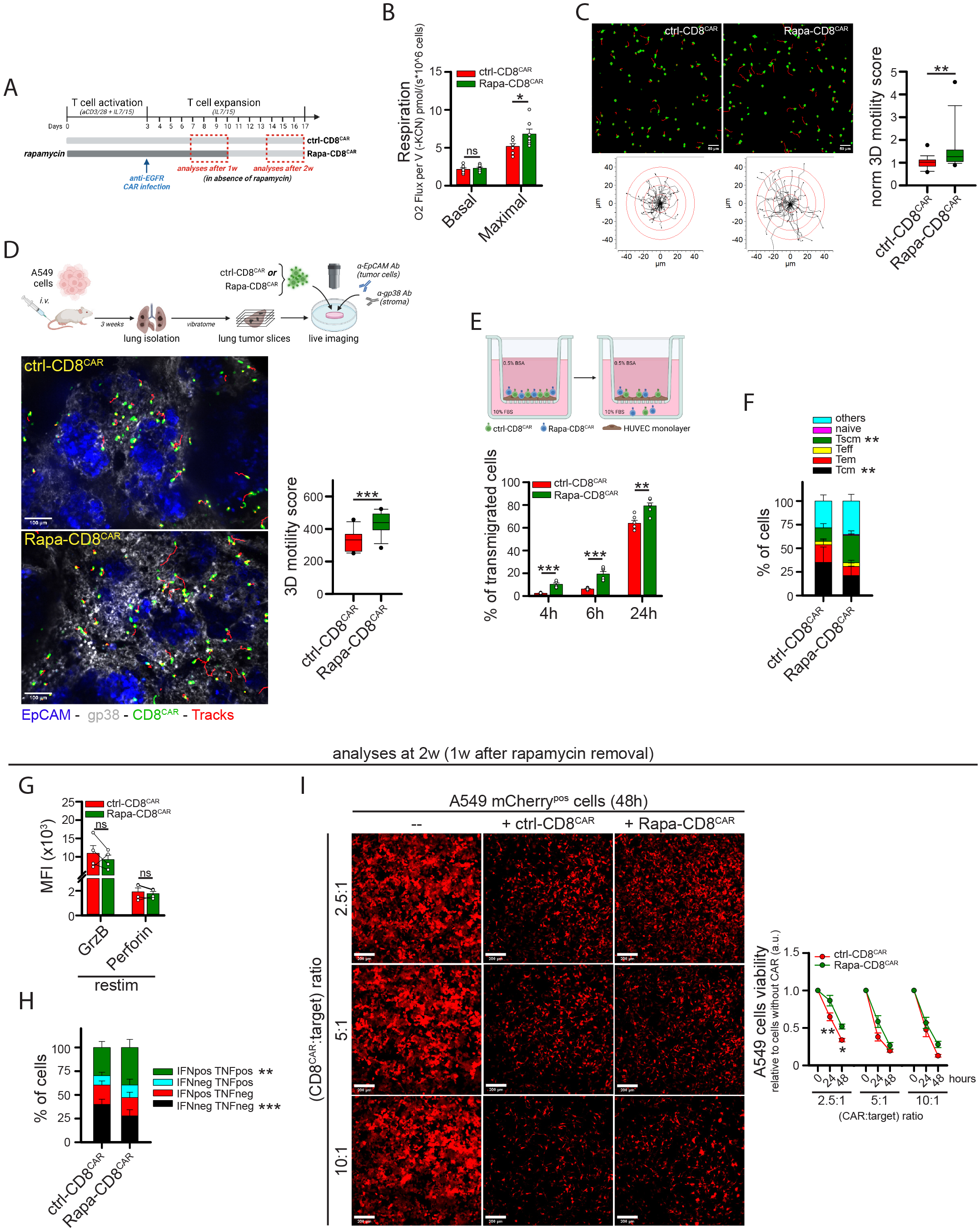
Rapamycin treatment promotes intra-tumoral motility and transmigration in anti-EGFR CD8^+^ CAR T cells. (**A**) Experimental layout. Anti-EGFR CD8^+^ CAR T cells were cultured in a control medium (ctrl-CD8^CAR^) or in the presence of 10nM rapamycin (Rapa-CD8^CAR^). All assays were performed in the absence of rapamycin. (**B-C**) Measurements of basal and maximal respiration (B, n=7) and normalized 3D motility Score (C, n=18). (**D**) Experimental layout on top. 3D motility of activated Calcein Green-stained ctrl-CD8^CAR^ and Rapa-CD8^CAR^ cells (cells in green, tracks in red) ovarlaid onto viable lung tumor slices (tumor islets in blue, EpCAM; stroma in gray, gp38) derived from i.v. injection of A549 lung tumor cells into NSG mice (n=12). (**E**) Percentage of Calcein Green-stained ctrl-CD8^CAR^ and eFluo670-stained Rapa-CD8^CAR^ cells (mixed at 1:1 ratio) that transmigrated across a TNFα-activated HUVEC monolayer in a transwell system (n=6). (**F**) Relative proportion of indicated T cell subsets in ctrl-CD8^CAR^ and Rapa-CD8^CAR^ cells (n=4). (**G-H**) At 2w (4/7d after rapamycin removal) cells were restimulated for 5h with human TransAct + BfdA. GranzymeB (GrzB) and Perforin levels (MFI) are indicated in G (n=4). For IFNγ and TNFα, the percentage of cells producing these cytokines is indicated in H (n=4). (**I**) Cytotoxicity of ctrl-CD8^CAR^ and Rapa-CD8^CAR^ cells against A549 (EGFR^+^ mCherry^+^) tumor cells. A549 cell viability was evaluated by measuring mCherry fluorescence (MFI) for each microscope field at the indicated time points (n=12). Scale bar, 50µm in C, 100µm in D and 200µm in I.

Besides migration, a functional CAR T cell must display additional activities, such as persistence in the host, cytotoxicity, and cytokine production. Therefore, we tested whether these functions were affected by rapamycin treatment. Consistent with what was observed for Rapa-CD8 cells, Rapa-CD8^CAR^ cells showed a more primitive (Tscm-like) state (**Fig. 6F**), which could presumably be responsible for the reduced *in vitro* expansion rate of these cells (***Fig. S9D***). Cytotoxicity and cytokine production were assessed 1-week after rapamycin removal (see scheme in **Fig. 6A**), as we reasoned that Rapa-CD8^CAR^ cells should only fulfill these functions once in the TME and therefore several days after injection into patients (and rapamycin removal). Compared with ctrl-CD8 ^CAR^, Rapa-CD8^CAR^ cells showed increased production of IFNγ and TNFα cytokines upon restimulation (**Fig. 6G-H**) and slightly reduced cytotoxicity against human A549 (EGFR^+^) lung tumor cells but only at a low CAR:target ratio (**Fig. 6I**).

In summary, rapamycin treatment generates functional CD8^+^ CAR T cells empowered with superior infiltration ability and intra-tumoral motility.

### Rapamycin-treated human CD8^+^ CAR T cells are more effective at infiltrating solid tumor masses in preclinical solid tumor xenograft models

We then decided to test whether Rapa-CD8^CAR^ cells also showed enhanced infiltration into solid tumor islets *in vivo*. To this aim, we chose two preclinical models based on the injection of human A549 lung tumor cells (EGFR^+^) into NSG mice. First, A549 cells were i.v. injected to generate lung tumors (orthotopic model, see scheme in **Fig. 7A**). After 3 weeks, mice were i.v. inoculated with anti-EGFR ctrl- or Rapa-CD8^CAR^ cells. To evaluate CAR T cell infiltration and ability to reach tumor islets, mice were sacrificed 4 days after, and lungs were analyzed. The total amount of ctrl- or Rapa-CD8^CAR^ cells infiltrating the lungs was similar (**Fig. 7B** and ***Fig. S10A***), consistent with the notion that the lungs are a particularly favorable site for T cell infiltration^48^. However, we observed that Rapa-CD8^CAR^ cells were significantly more accumulated in tumor islets compared with ctrl-CD8^CAR^ cells, which remained largely in the surrounding stroma (**Fig. 7B**). Of note, such an increased infiltration of Rapa-CD8^CAR^ cells was also associated with higher levels of active-caspase-3 (a marker of apoptosis) in tumor islets (**Fig. 7C**), indicating a stronger anti-tumor response in this model. We also assessed the ability of Rapa-CD8^CAR^ cells to reduce tumor growth in this model by injecting suboptimal amount of CAR T cells into NSG mice bearing a luciferase^pos^ A549 cells-derived lung tumor injected 3 days before (see scheme in **Fig. 7D**). Interestingly, mice inoculated with Rapa-CD8^CAR^ cells, but not with ctrl-CD8^CAR^, showed a significant reduction in tumor growth compared with untreated mice (**Fig. 7D**).

**Figure 7.**
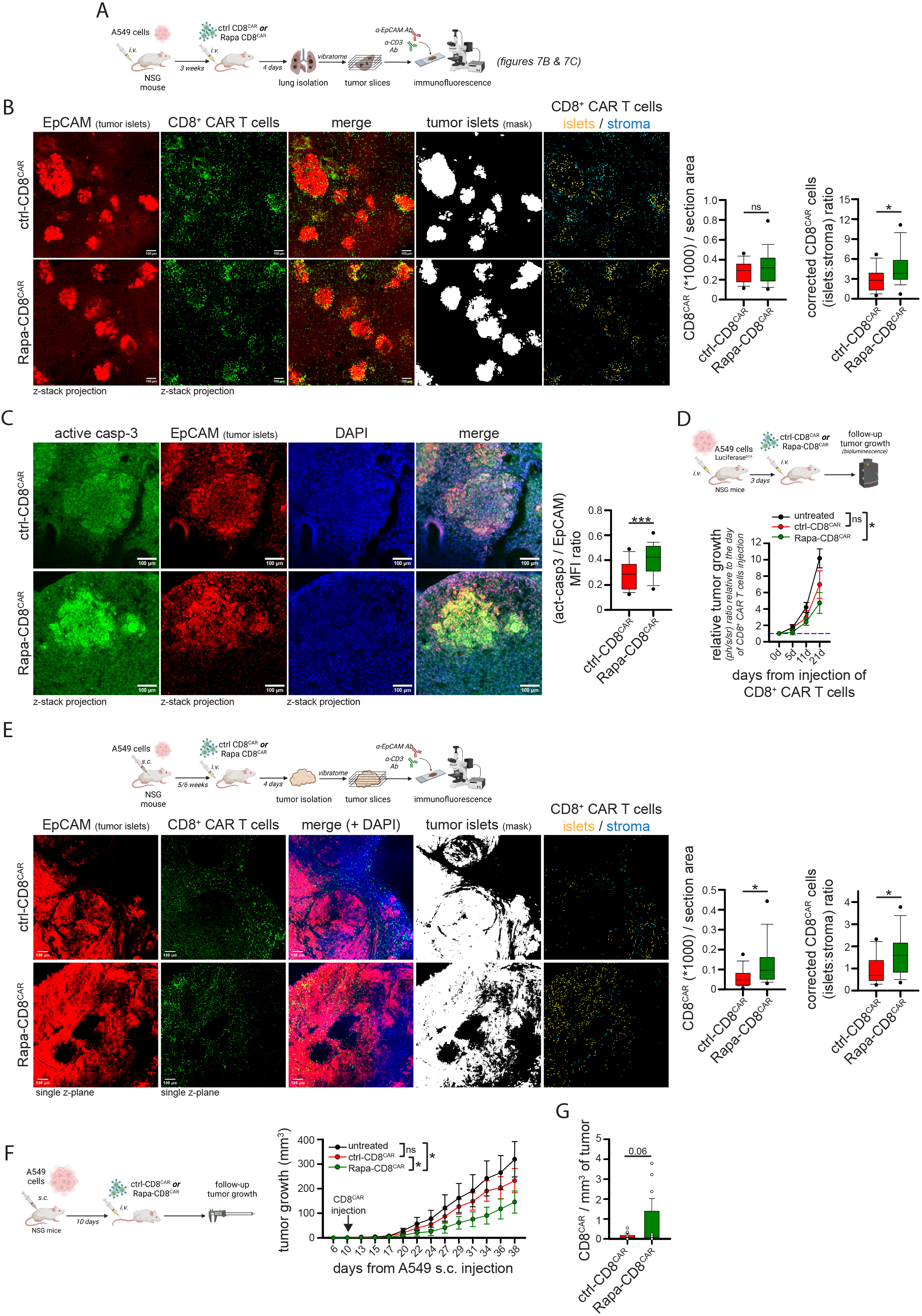
Rapamycin-treated anti-EGFR CD8^+^ CAR T cells show enhanced infiltration and better tumor control in two different xenograft tumor models. (**A-C**) Experimental layout in A. In B, representative immunofluorescence images of anti-EGFR ctrl-CD8^CAR^ or Rapa-CD8^CAR^ cells (anti-CD3 Ab, green) distribution in tumor islets and stroma (tumor cells identified using a EpCAM Ab, red) in each microscope field (ctrl-CD8^CAR^, n=17; Rapa-CD8^CAR^, n=27). An ImageJ mask was used to identify tumor islets (EpCAM^+^), stroma (EpCAM^neg^) and CAR T cells. Graphs on the right report the total amount of CAR T cells per section area (left) and the relative CAR T cell (tumor islets/stroma) ratio (corrected for the [tumor islets/stroma] area ratio). In C, immunofluorescence images of lung tumor slices stained for EpCAM (red) and active-caspase-3 (green). The graph on the right shows the relative active-caspase-3/EpCAM ratio for each tumor islet (ctrl-CD8^CAR^, n=136; Rapa-CD8^CAR^, n=102). (**D**) Experimental layout on top. Measurement of tumor size by bioluminescence (normalized to value at day of CAR T cell injection) in mice bearing lung tumors (derived from A549 i.v. injection 3 days before) untreated or infused with ctrl-CD8^CAR^ or Rapa-CD8^CAR^ cells (untreated, n=2; ctrl-CD8^CAR^, n=3; Rapa-CD8^CAR^, n=3). (**E**) Experimental layout on top. Images and graphs are the same as in B (ctrl-CD8^CAR^, n=18; Rapa-CD8^CAR^, n=14) with the exception that an anti-CD45 Ab was used to identify CAR T cells. (**F-G**) Experimental layout on left. Measurement of tumor size in A549-derived s.c. tumor-bearing mice untreated or infused with ctrl-CD8^CAR^ or Rapa-CD8^CAR^ cells (F; untreated, n=7; ctrl-CD8^CAR^, n=8; Rapa-CD8^CAR^, n=8). Amount of CD8^+^ CAR T cells per mg of tumor at endpoint (38 days) (G; ctrl-CD8^CAR^, n=8; Rapa-CD8^CAR^, n=7). Scale bar, 100µm in B-C-E.

To evaluate the effectiveness of Rapa-CD8^CAR^ cells in a different TME, A549 cells were s.c. inoculated into NSG mice to generate a subcutaneous tumor (see scheme in **Fig. 7E**). The morphology of TME is altered in this model compared with the orthotopic one (tumor cells do not accumulate in stroma-surrounded islets but in large areas with fewer adjacent stroma) and vasculature is more deregulated (as frequently observed in human solid tumors^49^), making CAR T cell infiltration more difficult compared with lungs^48^. Mice were i.v. inoculated with anti-EGFR ctrl- or Rapa-CD8^CAR^ cells 5/6 weeks after tumor cell inoculation, and then sacrificed 4 days after. Remarkably, despite no alterations in the amount of tumor cells per section (***Fig. S10B***), Rapa-CD8^CAR^ cells showed increased infiltration into the tumor mass, as indicated by an increased amount of CAR T cells per section (**Fig. 7E**), and also accumulated significantly more in contact with tumor cells compared to ctrl-CD8^CAR^ cells, which remained confined in the few stromal areas (**Fig. 7E**). To evaluate anti-tumor activity in this model, CD8^+^ CAR T cells were infused in tumor-bearing NSG mice 10 days after s.c. injection of A549 cells and tumor growth was followed for 1 month (see scheme in **Fig 7F**). Noteworthy, while injection of ctrl-CD8^CAR^ cells had no effect, Rapa-CD8^CAR^ cells significantly reduced tumor growth compared to untreated mice (**Fig. 7F**, see also ***Fig. S10C-D***). Last, we isolated tumor-infiltrating CD8^+^ CAR T cells at 38 days (*i.e.*, 1 month after CAR T cell injection) and we found an increased although not significant (p=0.06) number of Rapa-CD8^CAR^ cells in s.c. tumors compared with ctrl-CD8^CAR^ cells (**Fig. 7G**). We speculate that this increase may be associated with the more primitive (Tscm-like) initial differentiation state of Rapa-CD8^CAR^ cells (see **Fig. 6F**), which has been previously associated with improved CAR T cell long-term persistence^50–52^.

Taken together, our data demonstrate that *in vitro* rapamycin treatment generates CD8^+^ CAR T cells with enhanced ability to infiltrate into tumor islets, leading to a better control of tumor growth in two preclinical solid tumor models.

### G2Activation at 39°C improves mitochondrial metabolism and intra-tumoral motility in CD8^+^ CAR T cells

To reinforce the idea that manipulation of mitochondrial metabolism increases CD8^+^ CAR T cell 3D motility, we used an alternative drug-free system by taking advantage of a recent finding describing that culturing CD8^+^ T cells under fever-like conditions (*i.e.*, 39°C) increases mitochondrial activity^53^. CD8^+^ T cells were activated for 3 days at either 37°C (act-37°C) or at 39°C (act-39°C) and then transferred to 37°C for viral infection (anti-EGFR CAR) and subsequent expansion (see scheme in **Fig. 8A**). Activation at 39°C had no impact on cell viability (***Fig. S11A***) but increased maximal mitochondrial respiration (**Fig. 8B**) without affecting mitochondrial mass, MMP, mtROS (***Fig. S11B***), glycolysis (**Fig. 8C**), *in vitro* expansion (**Fig. 8D**) and cytokine production (***Fig. S11C***). Remarkably, CD8^+^ CAR T cells activated at 39°C showed (*i*) increased 3D motility in collagen gel (**Fig. 8E** and **Movies S53-56**), an effect likely promoted by the increased mitochondrial activity since oligomycin abrogated it (**Fig. 8E** and **Movies S53-56**), and (*ii*) superior intra-tumoral motility when deposed onto viable tumor slices derived from the BxPC3-NSG model (**Fig. 8F** and **Movies S57-58**).

**Figure 8.**
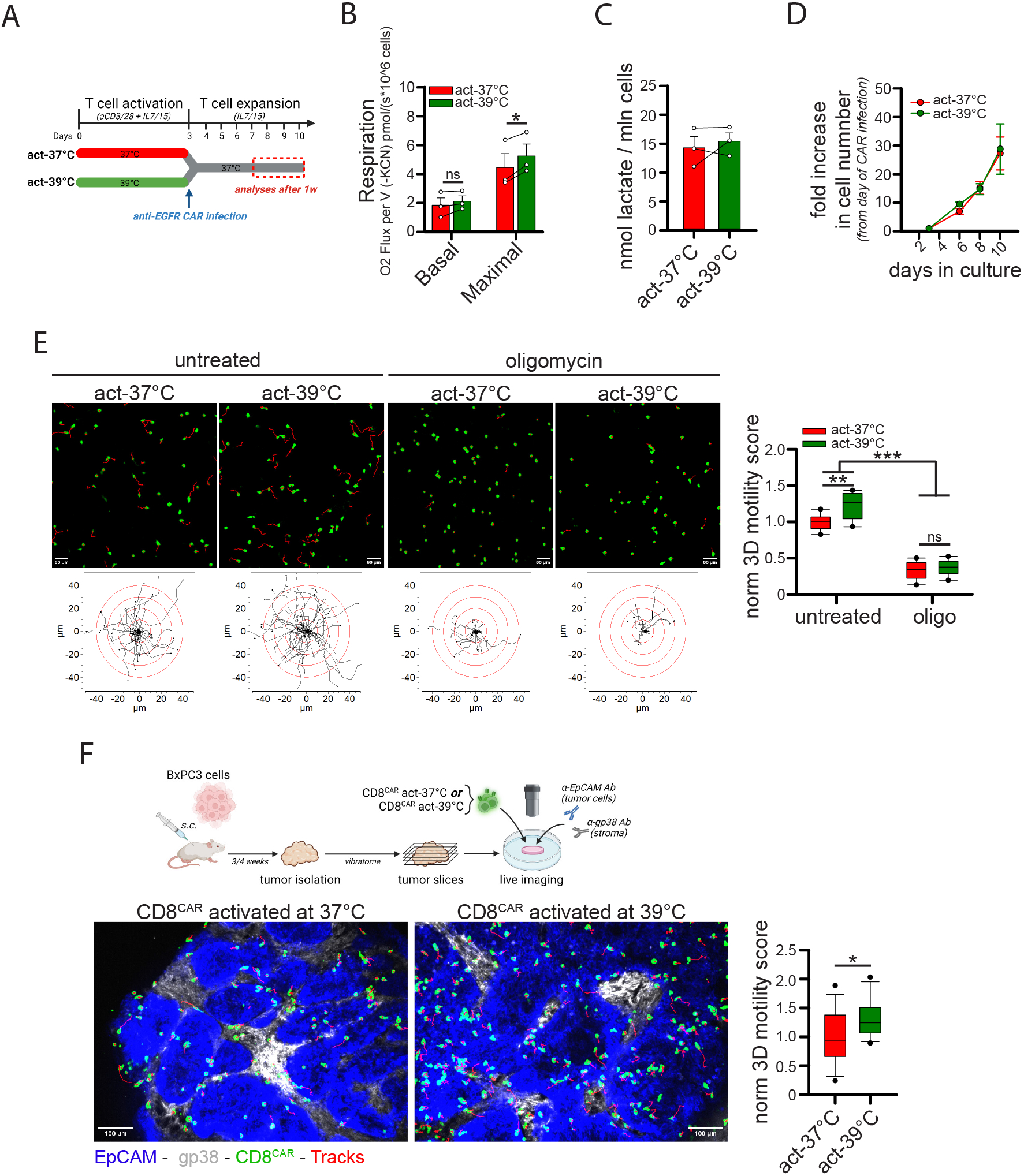
CD8^+^ CAR T cells activated at 39°C show higher mitochondrial respiration and better 3D intra-tumoral motility. (**A-E**) Experimental layout in A. CD8^+^ T cells activated at 37°C (act-37°C) or at 39°C (act-39°C) for 3d have been placed back at 37°C and infected with lentiviral particles to generate anti-EGFR CD8^+^ CAR T cells and were subsequently expanded at 37°C. After 1w, the following parameters were measured: basal and maximal respiration (B, n=3), lactate production (C, n=3), fold increase in cell number (D, n=3), and normalized 3D motility Score (E, n=9). (**F**) Experimental layout on top. 3D motility of Calcein Green-stained act-37°C or act-39°C cells (cells in green, tracks in red) overlaid onto viable tumor slices derived from BxPC3/NSG model (tumor islets in blue, EpCAM; stroma in gray, gp38) (n=16). Scale bar, 50µm in E and 100µm in F.

Overall, these data demonstrate that cell activation at 39°C can be an efficient strategy to increase mitochondrial activity and hence intra-tumoral motility of CD8^+^ CAR T cells.

## DISCUSSION

The ability of T cells to migrate and reach tumor cells determines the efficacy of T cell-based immunotherapy. Thus, there is a lot of interest in elucidating the mechanisms responsible for the migration of CD8 T cells in tumors and in developing strategies to boost this function. Studies performed using imaging techniques have underlined the importance of both environmental and cell-intrinsic factors in this process. In the present work, we identified the metabolic requirements of CD8^+^ T cell migration in a 3D environment (especially intra-tumoral one). Our results obtained in vitro and confirmed in ex vivo tumor slices indicated a key role of the TCA cycle fueled by glucose and glutamine, but no FAs, with glycolysis playing only a much minor role (***Figure S12***). Strategies targeting mitochondrial metabolism were then developed and tested in several preclinical models including two mouse tumor models. We demonstrated that CD8^+^ CAR T cells with enhanced mitochondrial activity showed superior infiltration into solid tumors leading to better inhibition of tumor growth (***Figure S12***).

The poor role of glycolysis in supporting the amoeboid-like of CD8^+^ T cells in 3D environments contrasts clearly with what has been observed for migrating cancer cells. Here, glycolysis plays a much more important role by providing ATP locally at lamellipodia to sustain formation of protrusion^54,55^. Moreover, mtROS have been described to positively support cancer cell migration and metastatic spread^56^. Interestingly, rotenone treatment has been shown to promote cancer cell dissemination at doses that would inhibit OXPHOS-derived ATP generation^56^, further reinforcing the role of glycolysis for energy production in these migrating cells. We also observed a positive role for mtROS in CD8^+^ T cell 3D motility. However, given the prominent role of mitochondrial metabolism and not glycolysis, a subtle balance between mtROS and mitochondrial ATP (mito-ATP) production is required in CD8^+^ T cells to support 3D motility, since an excessive increase in mtROS amount would be detrimental for their migration because of the concomitant reduction in mito-ATP production. This is consistent with several studies showing that high ROS levels inhibit amoeboid-like motility^32^. Our data are also in line with previous reports suggesting that mito-ATP is required to support T cell motility in a 2D environment by providing energy for the acto-myosin contraction at the cell rear edge^13,16,57^. Low amounts of ROS, by modulating the redox status of many proteins, are positive contributors to T cell activation^58,59^. The exploration of the nature of the molecular targets of mtROS that control CD8^+^ T cell motility will be an important area for the future.

Several strategies modulating mitochondrial metabolism proved to be effective in increasing the efficacy of T and CAR T cell-based immune-therapy approaches^21,60–63^. The explanation to justify these effects relies on the ability of these treatments to increase T cell differentiation towards more memory/primitive-like states with superior long-term persistence^21,61,64,65^. However, the impact of these strategies on modulating T cell intra-tumoral motility or T cell infiltration into tumor islets has never been addressed in these studies. Based on our findings, we propose that along with an improved memory T cell differentiation, the beneficial effects of these treatments can also be explained by an enhanced migration of T cells in tumors favoring the formation of productive contacts with cancer cells. In support of this, the combination of IL-7 and IL-15 cytokines, which are commonly used during CAR T cell production, promotes both memory T cell expansion and intratumoral T cell migration.

From a clinical perspective, we here propose to pharmacologically manipulate CD8^+^ CAR T cell metabolism during the *in vitro* expansion phase and prior to their injection into patients to confer on these cells a superior and long-lasting motility capacity while at the same time avoiding any possible alterations on tumor cell metabolism, which could unintentionally promote tumor growth. Alternative approaches to increase mitochondrial metabolism in CD8^+^ CAR T cells also consist of the genetic engineering of lymphocytes to overexpress metabolic enzymes^66^. However, we believe that the rapamycin-based strategy that we tested here would be preferable to genetic approaches permanently increasing mitochondrial activity, which would not allow T cells to engage different metabolic pathways when required. For example, during phases of cell proliferation, the OXPHOS rate must be reduced and ATP must be mainly produced by glycolysis to spare carbons for biosynthesis^26^. In this context, forcing T cells to permanently maintain a high OXPHOS rate could be detrimental. On the contrary, the rapamycin-based strategy did not alter basal respiration levels but only maximal respiratory capacity, indicating that CD8^+^ T cells maintain the flexibility to increase OXPHOS only when required (such as during migratory phases), while reducing it when needed (such as during proliferative phases). Unfortunately, given its role in inhibiting mTOR-dependent glycolysis, rapamycin treatment reduced the *in vitro* expansion rate of CD8^+^ CAR T cells. Although this may not be a limitation for most cancer patients, it is possible that this reduced proliferation rate would be an obstacle to generate enough CAR T cells from immunocompromised patients lacking a high initial quantity of T cells. In this case, alternative approaches not affecting proliferation could be preferable. For example, we tested bezafibrate treatment, which had no impact on cell proliferation, although its effects on 3D motility were less pronounced and no increase in mitochondrial mass was observed (therefore limiting its long-lasting efficacy). Further studies will be required to implement the best metabolic reprogramming strategies enabling T cells to migrate actively, expand and survive within the tumor microenvironment.

Hypoxia is a major obstacle to anti-cancer approaches. Besides supporting cancer cell metabolic reprogramming and the Warburg effect^67,68^, hypoxia also prevents T cells from mounting a potent antitumor response^69^. In this context, our data suggesting the prominent role of mitochondrial metabolism in supporting CD8^+^ T cell intra-tumoral motility are in line with the observation of a strong reduction in T cell motility within solid tumor hypoxic areas^70^. Therefore, it is possible that strategies aimed at increasing mitochondrial activity to support T cell intra-tumoral motility would not produce the expected results in a highly hypoxic TME devoid of O_2_. In this case, alternative strategies should be envisaged. For example, culturing cells in mild acidosis increases ETC efficiency, allowing efficient OXPHOS even at low oxygen tension^71^. It would be interesting to test this strategy in a context of CAR T cells.

Metabolic alterations within the TME are known to impact T cell activities^72^. Our data suggest that long-time glucose or glutamine deprivation in the TME may negatively influence the ability of endogenous or CAR CD8^+^ T cells to properly infiltrate tumor islets. Surprisingly, FAs do not appear to be involved in supporting this process, at least when provided as directly available energy sources to migrating CD8^+^ T cells. Although metabolic strategies to sustain or rescue FAO proved to be effective in increasing TILs functions^45,63^, these are mainly long-term treatments, presumably reprogramming metabolism to reinforce mitochondrial activity. In this regard, it would be interesting to test whether long-term FAs supplementation could have any effect on CD8^+^ T cell 3D motility. In addition, our data indicate that lactate can be used by CD8^+^ T cells to sustain intra-tumoral motility. Therefore, although lactate has been reported to play an inhibitory role for CD8^+^ T cells in the TME, especially for proliferation and cytotoxicity^73,74^, our data suggest that its role may be more multifaceted, at least supporting motility in the absence of glucose. Of note, regulatory T cells (Tregs) can use lactate in the TME to sustain their OXPHOS metabolism, providing them some advantage over conventional T cells (Tconvs)^75^. No study to date has addressed the relative distribution of Tregs and Tconvs within solid TME. Based on our results, it would be interesting to investigate whether lactate might enable Tregs to better infiltrate tumor masses compared to Tconvs cells (especially in a glucose-deprived TME) as a mechanism to support tumor growth.

In conclusion, we identified here the metabolic requirements of CD8^+^ T cells migrating in 3D environments, such as within a solid tumor mass. Our study provides a rationale for manipulating metabolism of CD8^+^ CAR T cells to improve infiltration of these cells into tumor islets and potentiate their anti-tumor effect.

## MATERIALS & METHODS

### Study approvals

Human studies were carried out according to French law on biomedical research and to principles outlined in the 1975 Helsinki Declaration and its modification. Peripheral blood samples were obtained from EFS (Etablissement Français du Sang) under agreement 18/EFS/030, including informed consensus for research purposes. Samples were anonymous and not associated with any personal data. Human lung biopsies were obtained from the Pathology Service of Cochin Hospital. Ethical procedures were approved by CPP Ile de France II (approval number 00001072, August 27, 2012) and INSERM Institutional Review Board (CEEI, IRB00003888, approval number 22/893, March 8, 2022), including informed consensus for research purposes.

NOD.Cg-Prkdc(scid)Il2rg(tm1Wjll)/SzJ (abbreviated NSG) immunodeficient mice (The Jackson Laboratory, Bar Harbor, ME, USA, 005557) were housed in pathogen-free condition at Cochin Institute Animal Facility. The mice protocol has been approved by Paris Descartes University (CEEA 17-039) and the French Ministry of Research (APAFiS #15076).

### Cell culture and reagents

Human peripheral blood mononuclear cells (PBMCs) were isolated using Ficoll gradient (800g for 20min, RT) and T cells were purified using Human Naive CD8^+^ T Cell Isolation Kit (Miltenyi, 130-093-244), Naïve CD4^+^ T cell Isolation Kit (Miltenyi, 130-094-131) or Pan T cell Isolation Kit (Miltenyi, 130-096-535). Then T cells have been collected, washed, and used for subsequent analyses. Human T cells have been cultured in RPMI 1640 medium (Thermo Fisher 21875) supplemented with 10% Fetal Bovine Serum (Thermo Fisher A3840002), 2mM L-glutamine (Thermo Fisher 25030081) and 100U/ml penicillin/streptomycin (Thermo Fisher 15140130). When indicated, RPMI medium glucose-(Thermo Fisher 11879) or glutamine-free (Thermo Fisher 31870) was used.

For *in vitro* activation, 8×10^5^ naïve CD8^+^ or CD4^+^ T cells have been stimulated with 10µg/ml anti-CD3 (plate-coated) (Biolegend 300465) and 2µg/ml anti-CD28 (Biolegend 302943) for 72h in 96well plate in presence of 25ng/ml IL-7 and IL-15 (Miltenyi, 130-095-363 and 130-095-765), or in the presence of 200U/ml IL-2 alone (Miltenyi, 130-097-746). Then, cells have been washed and expanded with ILs only. When indicated, cells were cultured in the presence of 10nM rapamycin (Tocris 1292), 10µM bezafibrate (Sigma Aldrich B7273), 1mM AICAR (Tocris 2840) or 1mM βNAD (Sigma Aldrich N1511). Drugs were removed on the day of assays and cells were washed at least twice with RPMI complete medium. Unless specified, these drugs were not present during the assays.

To evaluate T cell motility in collagen gels or viable tumor slices, T cells were resuspended in a “*motility medium*” composed of RPMI 1640 medium supplemented with 2mM glutamine, 0.5% Bovine Serum Albumin (BSA) (Sigma Aldrich A3059) and 10mM HEPES (Thermo 15630). When specified, a glucose- or glutamine-free RPMI medium was used to prepare the *motility medium*.

The following reagents were added either during motility assay or in cell culture for the indicated period of time before the assay: 11mM glucose (Thermo Fisher A2494001), 2mM or 11mM glutamine, 85µg/L Oleic Acid (BSA-conjugated, Aigma Aldrich O3008), 130-13000µg/L Palmitate (BSA-conjugated, CaymanChemical 29588), 85-8500µg/L Linoleic Acid (BSA-conjugated, Sigma Aldrich L9530), 5µM etomoxir (Sigma Aldrich 236020), 10µM trimetazidine (Sigma Aldrich 653322), 1µM perhexiline (Sigma Aldrich, SML0120), 20mM 2-deoxy-glucose (2DG) (Sigma Aldrich D8375), 100µM 6,8-bis(benzylthio)octanoic acid (indicated as OctanoicAcid, OA, Sigma Aldrich SML0404), 10µM CB-839 (Euromedex TA-T6797), 11mM galactose (Sigma Aldrich PHR1206), 10mM sodium lactate (Sigma Aldrich L7022), 1mM sodium acetate (Sigma Aldrich 71188), 100µM sodium pyruvate (Thermo Fisher 11360070), oligomycin (several doses, Sigma Aldrich 495455), rotenone (several doses, Calbiochem B48409), antimycin-A (several doses, Sigma Aldrich A8674), 3-nitropropionic acid (3-NP, several doses, Sigma Aldrich N5636), Atpenin-A5 (several doses, MedChemExpress HY-126653), MitoTEMPO (several doses, Sigma Aldrich SML0737), 500µM N-acetyl-cysteine (NAC, Sigma Aldrich, A9165), 1µM PS-10 (Sigma Aldrich SML2418), 1µM LDH inhibitor GSK2837808A (LDHi, Tocris 5189), 10µM α-Lipoic Acid (Sigma Aldrich 62320), 1mM succinyl phosphonate (MedChemExpress HY-12688A), 100µM di-methyl-succinate (Sigma Adrich 112755), 100µM di-methyl-α-ketoglutarate (Sigma Aldrich 349631), 500nM CCCP (Alfa Aesar L06932.ME), and 50nM ethidium bromide (Sigma Aldrich E1385).

Human BxPC3 tumor cells (in-house stock) have been cultured in the same medium as T cells. Human A549 (mCherry^pos^ and Luciferase^pos^) and HEK-293T tumor cells (in-house stocks) have been cultured in DMEM medium (Thermo Fisher 11995) supplemented like RPMI medium for T cells. HUVEC cells (Thermo Fisher C0035C) have been cultured in Human Large Vessel Endothelial Cell Basal medium (Thermo Fisher M200500) supplemented with 10% FBS and Large Vessel Endothelial Supplement (Thermo Fisher A1460801). Cells were routinely tested for mycoplasma contamination.

### In vitro motility assay with collagen gel

T cells were washed with DPBS, stained with 100nM of Calcein Green (Invitrogen C34852) or Calcein Red Orange (Invitrogen C34851) for 20min at 37°C in DPBS, washed once in motility medium, resuspended in 50µl at a final concentration of 20×10^6^cells/ml and then mixed with 100µl of collagen solution (PureCol Bovine Collagen, CellSystem 5005, previously neutralized using DPBS 10x) to reach a final collagen concentration of 2mg/ml. Then, the mixture was loaded into an ibiTreat μ-Slide VI 0.4 slide (Ibidi 80606), which was placed at 37°C in the incubator for 1h to allow collagen polymerization. Motility was always evaluated in the absence of any interleukin. Drugs or specific nutrients were added to the medium immediately before mixing cells and collagen. When indicated, telo-collagen (TeloCol purified Bovine Collagen, Cellsystem 5026) was used at a final concentration of 3mg/ml. Live recording of T cell motility was performed using a Nikon epifluorescence microscope equipped with 37°C heated-chamber, ORCA-Flash 4.0 camera (C11440 Hamamatsu) and Metamorph software (version 7.10.2.240, Molecular Devices). For each condition, 15 images (30sec time interval, 4/5 z-planes, z-distance 15µm) were acquired using a 20x objective (binning rate, 2) at 3 different stages. In most cases, T cells from two different donors were stained with Calcein Green and Calcein Red and mixed before adding them to collagen. In other experiments, cells from different experimental conditions but from the same donor were stained with Calcein Green and Calcein Red and then mixed before adding collagen.

### Live imaging slice assay

Viable tumor slices were obtained from human NSCLC biopsies, BxPC3- or A549-derived tumors implanted subcutaneously into NSG mice or lungs of NSG mice inoculated i.v. with A549 cells (orthotopic lung tumor model). Slices were obtained and prepared essentially as previously described^76^. Briefly, tumor samples were cut into 1-2mm^3^ pieces, embedded into 7% agarose (Sigma Aldrich A0701) and cut into through vibratome (VT 1000S, Leica) to obtain 300µm-thick viable tumor slices (Leica Biosystem). Then, slices were stained with anti-EpCAM (BV421 or PE, Biolegend 324220 or 324206) and anti-gp38 (eFluor660, eBioscience 50-538182 or BV421, Biolegend 127423) Abs using Millicell 0.4µm organotypic insert (Millipore PICMORG50) and metal rings for 30min at 37°C. NSCLC biopsies were also stained with PE-anti-CD8 Ab (Biolegend 344706) to detect endogenous CD8^+^ T cells and Alexa647-anti-fibronectin Ab (BD Bioscience 563098) to detect stroma (instead of gp38). When exogenous T cells were added to the slice, T cells stained with Calcein Green and/or Calcein Red (see above) were either (*i*) added onto the stained slice (5×10^5^ cells/30µl of motility medium) for 20min at 37°C or (*ii*) injected into the tumor piece with 30G syringe (10×10^6^ cells/100µl of motility medium) before embedding the tumor biopsy into agarose. Finally, slices were placed into a 30mm petri dish (Thermo Fisher 121V) filled with 5ml of motility medium and let warm at 37°C for 20min in the incubator. Live recording of cell motility was performed using a spinning-disk (Yokogawa CSU-X1) Leica DM6000FS microscope equipped with 37°C heated-chamber, ORCA-Flash 4.0 camera (C11440 Hamamatsu) and Metamorph software (version 7.8.9, Molecular Devices). For each condition, 15/20 images (5/10 z-planes, z-distance 15µm) were acquired using a 10x or 25x objective (binning rate, 2) with a time interval of 30sec.

### Analysis of T cell motility

T cell motility was analyzed using ImageJ plugin TrackMate^77^ in a 3D environment (z-stack distance, 15µm). LoG detector was used to identify spots (quality and maximal intensity filters applied for collagen assays) and Simple LAP Tracker to identify tracks (linking max distance 20µm, closing-gap max distance 20µm, frame gap 3). Only tracks containing 10 or more spots were considered for analysis. Motility Score was calculated as follows: (% of cells moving more than 10µm as Euclidean distance)x(mean speed µm/min of moving cells). Images and movies shown throughout the manuscript are 2D z-stack reconstruction while analyses were performed on original 3D images (also considering z-axis displacement).

### Immunofluorescence on fixed samples

Samples were fixed o.n. with PLP solution (1% paraformaldehyde, 2mM lysine-HCl, 550mg/L NaIO_4_) at 4°C, cut into 1-2mm^3^ pieces, embedded into 7% agarose (Sigma Aldrich A0701) and then cut into 150µm-thick sections through vibratome (VT 1000S, Leica).

For extracellular staining, slices were incubated 1h at RT with 0.5% BSA in DPBS. For intracellular staining, slices were incubated for 20min at −20°C with 100% cold acetone (Carlo Erba 400971), washed twice with DPBS, and incubated for 1h at RT with 1% BSA + 0.3% TritonX-100 (Sigma Aldrich T9284) in DPBS. Then, slices were placed onto Millicell 0.4µm organotypic insert (Millipore PICMORG50) and 30µl of 0.5% BSA / DPBS solution (for extracellular staining) or 0.5% BSA + 0.1% Tween20 (Sigma Aldrich P7949) / DPBS solution (intracellular staining) containing the desired Abs were added. A metal ring was used to keep the solution in place on the slice. Abs were left incubated o.n. at 4°C. The day after, slices were washed twice in DPBS and then mounted using Superfrost microscope slides (Epredia J1800AMNZ) and Vectashield antifade mounting medium (VectorLabs H-1900). The following antibodies were used: Alexa555-anti-TOMM20 (Abcam ab221292), Alexa647-anti-ATP5a (Abcam ab196198), PE-anti-IDH2 (Abcam ab212122), Alexa488-anti-HK1 (Abcam ab184818), Alexa647-anti-PFK2 (Abcam ab203651), Alexa405-anti-GLUT1 (Abcam ab210438), anti-G6PD (BioTechne NBP2-22125V), BV605-anti-EpCAM (Biolegend 324224), PerCP-Cy5.5-anti-CD8 (Biolegend 344710), Alexa647-anti-CD8 (Biolegend 344726), FITC-anti-CD3 (Biolegend 344804), PE-anti-EpCAM (Biolegend 324206), rabbit anti-cleaved caspase-3 (Cell Signaling 9664S) and Alexa647-anti-rabbit secondary antibody (Thermo Fisher A-21245). Images were acquired using a spinning-disk (Yokogawa CSU-W1 T1) Ixplore IX83 microscope (Olympus) equipped with ORCA-Flash 4.0 V3 (sCMOS Hamamatsu) camera and CellSens software (Olympus). To analyze protein expression in NSCLC samples, an ImageJ threshold mask was used to automatically identify EpCAM^+^ areas (tumor islets) and CD8^+^ T cells. Then, “subtract” or “multiply” ImageJ commands were used to define CD8^+^ T cells in tumor stroma or islets, respectively. Protein expression (MFI) was calculated in the selected T cell populations.

### CRISPR-Cas9 gene editing

CRISPR/Cas9-mediated gene editing was performed on CD8^+^ T cells activated for 24h. Briefly, 25pmol of each specific crRNA was mixed with an equal amount of tracRNA (IDT technologies 1072534) and warmed at 95°C for 5min. Then, the solution was left cool down to RT and 15pmol of Cas9 Nuclease V3 (IDT Technologies 1081059) were added. After 20min incubation at RT, 16.4µl P3 and 3.6µl Supplement Reaction solutions from P3 Primary Cell 4D X Kit S (Lonza V4XP-3032) were added. Then, 5×10^5^ T cells were washed once in DPBS and resuspended in 20µl of the reaction mixture. Electroporation was performed using Amaxa 4D Nucleofector (Lonza), following the manufacturer instruction. After 30min recovering at 37°C, cells were restimulated for an additional 48h. Target sequences (from IDT Technologies) used for gene editing were the following (three sequences were used together to target PDHA1 and/or OGDH):

5’-CACGGCTTTACTTTCACCCGGGG-3’ (Hs.Cas9.PDHA1.1.AA),

5’-TATGCCAAGAACTTCTACGGGGG-3’ (Hs.Cas9.PDHA1.1.AB),

5’-TAGAGCAATCCCAGCGCCCAGGG-3’ (Hs.Cas9.PDHA1.1.AC)

5’-TCCACGAGCTTGTCCACGTTGGG-3’ (Hs.Cas9.OGDH.1.AA)

5’-TGGGACTAGTTCGAACTATGTGG-3’ (Hs.Cas9.OGDH.1.AB)

5’-ACGAATGCCGGAGCCCCACCGGG-3’ (Hs.Cas9.OGDH.1.AC)

### qRT-PCR for mitochondrial DNA

DNA extraction was performed with FastPure Blood/Cell/Tissue/Bacteria DNA Isolation Mini Kit (Vazyme Biotech DC112-01). Cells were lysed in the presence of proteinase K (Sigma Aldrich 31158) at 56°C, then DNA was isolated by silica gel columns purification technology. Quantitative real-time PCR was performed with LightCycler 480 SYBR Green I Master detection kit (Roche 04707516001) on a Light Cycler 480 machine (Roche). Three technical replicates were performed for each sample. Fluorescence curve analysis was performed using LightCycler 480 Software (version 1.5.1.62, Roche). Primer sequences were the following:

ND1-Fw: 5’-CCCTACTTCTAACCTCCCTGTTCTTAT-3’

ND1-Rv: 5’-CATAGGAGGTGTATGAGTTGGTCGTA-3’

ND5-Fw: 5’-ATTTTATTTCTCCAACATACTCGGATT-3’

ND5-Rv: 5’-GGGCAGGTTTTGGCTCGTA-3’

18S-Fw: 5’-AGTCGGAGGTTCGAAGACGAT-3’

18S-Rv: 5’-GCGGGTCATGGGAATAACG-3’

### Lactate assay

0.5-2×10^6^ cells have been washed and resuspended in 500µl of *motility medium* in 96well plate for 4h. Then, cells have been collected and lactate has been measured by using Lactate Assay Kit (Sigma MAK064), following manufacturer instructions. Measurements were performed using CLARIOstar instrument (BMG Labtech).

### High-resolution respirometry with Oroboros device

The oxygen consumption rate (OCR) was measured in the *motility medium* (see above) by a high-resolution 2 mL glass chamber respirometer (Oroboros Oxygraph-2k). The electrode was calibrated at 37°C, 100% and 0% oxygen before adding CD8^+^ T cells (2mL at 2.5×10^6^ cells /mL) to each chamber and the flux of O_2_ consumption was measured in pmol/s. Oligomycin (0.5μg/mL) was added to block complex V, as an estimate of the contribution of mitochondrial leakage to overall cellular respiration. Increasing amounts of carbonyl cyanide m-chlorophenyl hydrazone (CCCP) were added to measure maximum respiratory capacity. Finally, potassium cyanide (KCN) was added to measure the non-mitochondrial respiration.

### Measurement of CO_2_ flux

15×10^6^ CD8^+^ T cells were incubated in RPMI 1640 medium (Thermo Fisher 31870) containing 11 mM [U-^14^C]-glucose (0.1mCi/mmol) or in RPMI 1640 medium (Thermo Fisher 11879) containing 2 mM [U-^14^C]-glutamine (0.1mCi/mmol). After 6h, the media were transferred to a conical glass vial for CO_2_ production measurement. Perchloric acid was injected into the incubation media through the rubber cap to a final concentration of 4% (v/v). Benzethonium hydroxide was injected through the rubber cap into a plastic well suspended above the incubation media. During 1 hour of vigorous shaking at 25°C, the released [^14^C]-CO_2_ was trapped by the benzethonium hydroxide and assessed by scintillation counting.

### Flow cytometry

The following antibodies have been used to stain extracellular proteins: BV711 anti-CD45RA (Biolegend 304138), APC anti-CD45RO (Biolegend 304210), PE anti-CCR7 (Biolegend 353204), APC-Cy7 anti-CD27 (Biolegend 302816), PE-Cy7 anti-CD95 (Biolegend 305621). True-Nuclear™ Transcription Factor Buffer Set (Biolegend 424401) was used to stain intracellular proteins, detected with the following anti-human antibodies: PE-anti-GrzB (Biolegend 372208), PE-Cy7-anti-Perforin (Biolegend 353315), APC-anti-IFNγ (Biolegend 502511), BV421-anti-TNFα (Biolegend 502932), Alexa647-anti-ATP5a (Abcam ab196198), PE-anti-IDH2 (Abcam ab212122), Alexa488-anti-HK1 (Abcam ab184818), Alexa647-anti-PFK2 (Abcam ab203651), anti-MCT1 (BioTechne FAB8275T), Alexa405-anti-GLUT1 (Abcam ab210438), anti-G6PD (BioTechne NBP2-22125V). Primary antibodies were incubated o.n. at 4°C. To evaluate GranzymeB, Perforin, IFNγ and TNFα production, 1×10^6^ cells have been restimulated for 5h in a 96well plate with 3µl of TransAct (Miltenyi 130-111-160). 5µg/ml BrefeldinA (Biolegend 420601) has been added for the last 2h and a half.

For T cell subpopulations, the following subsets were defined by flow cytometry gating strategy:

-) stem cell memory-like (Tscm) CD45RO^neg^CD45RA^pos^CD27^pos^CCR7^pos^CD95^pos^;

-) naïve-like (naive) CD45RO^neg^CD45RA^pos^CD27^pos^CCR7^pos^CD95^neg^;

-) effector-like (Teff) CD45RO^neg^CD45RA^pos^CD27^neg^CCR7^neg^;

-) central memory-like (Tcm) CD45RO^pos^CD45RA^neg^CD27^pos^CCR7^pos^;

-) effector memory-like (Tem) CD45RO^pos^CD45RA^neg^CD27^neg^CCR7^neg^.

Cells outside the indicated gates were defined as “others”.

To evaluate glucose uptake, mitochondrial membrane potential (MMP), mitochondrial mass or mitochondrial ROS (mtROS), cells were incubated for 20/30min at 37°C in the presence of 30μM 2-NBDG (Thermo N13195), 100nM TMRE (Thermo Fisher T669), 100nM MitoTrackerGreen (Thermo Fisher M7514) or 2.5μM MitoSox (Thermo Fisher 11579096), respectively. Then, cells have been washed and analysed. Mitochondrial mass (MitoMass) has been calculated as (MitoTrackerGreen/FSC) ratio. MMP has been calculated as (TMRE/MitoTrackerGreen) ratio, mtROS as (MitoSox/FSC) ratio.

Samples were analysed using BD Accuri C6 (CFlow Plus software) or BD Fortessa (DIVA software).

### Infiltration into collagen gel

5×10^5^ human BxPC3 tumor cells were resuspended into 450μl of complete RPMI medium plus neutralized collagen solution (2mg/ml) and inserted into one side chamber of μ-slide Chemotaxis Chamber (Ibidi 80326). The other side chamber was filled with medium alone. After 24h, T cells were stained with Calcein Green or Calcein Red as described above, mixed at 1:1 ratio and 2×10^6^ cells were resuspended into 80μl and inserted into the middle chamber. Z-stack images were acquired at 0h and 2h. ImageJ was used to identify cells and starting area (as ROIs) and to calculate the minimum distance.

### Anti-EGFR CAR and PercevalHR lentiviral infection

To generate anti-EGFR CAR, downstream of a signal peptide scFv sequences derived from nimotuzumab Ab (IMGT/2Dstructure-DB INN 8545H, 8545L) were introduced into the expression cassette encoding CD8 hinge and transmembrane domains (194-248 Aa, GenBank: AAH25715.1), 4-1BB costimulatory domain (214-255 Aa, GenBank: AAX42660.1), CD3z signaling domain (52-163 Aa, GenBank: NP_000725.1) and GFP coding sequence (downstream of IRES sequence). Production of lentiviral particles was performed by transient co-transfection (polyethylenimine:DNA at 3.5:1 ratio) of HEK 293T cells with the second-generation packaging system plasmid psPAX2 and pMD2.G (Addgene, 12260 and 12259). Viral supernatants were harvested at 48h post-transfection. Viral particles were concentrated by ultracentrifugation at 25.000g for 2h and conserved at −80°C. Anti-EGFR CD8^+^ T cells (defined as CD8^CAR^) were generated by infecting 6×10^5^ CD8^+^ T cells activated for 72h with 50µL (MOI2) of lentiviral particles in 600µL of RPMI complete medium (supplemented with IL-7+IL-15) in 48well plate. After 3 days, cells have been collected, washed, and used for subsequent analysis.

PercevalHR plasmid^38,39^ (Addgene 49083) was used to generate lentiviral particles using the same procedure as for “CAR” particles. After 1 week of stimulation, cells were stained with Calcein Red and motility was analysed in 2mg/ml collagen gel. PercevalHR and Calcein Red signals was measured at timepoint 0 and then the ratio was calculated. Cells were defined as motile or immotile by visual inspection following motility analysis of the TrackMate software.

### Cytotoxicity assay

Anti-EGFR CAR T cell cytotoxicity has been evaluated using mCherry^pos^ A549 cells. Briefly, A549 cells were plated at 1×10^5^ cells/250µl on µ-Slide 8 Well slides (Ibidi 80826) in DMEM medium. After 24h (time 0), anti-EGFR CAR T cells were added to obtain a CAR:target ratio of 2.5:1, 5:1 or 10:1 in RPMI complete medium. A549 cell viability was evaluated by quantifying mCherry fluorescence (MFI) at different time points with a fluorescent microscope and normalized to MFI values in absence of CAR T cells.

### Orthotopic and subcutaneous tumor models

For the orthotopic tumor model, 2×10^6^ human A549 cells were injected i.v. into the tail vein of 2 months-old NSG nude mice. After 3 weeks, mice were either (*i*) sacrificed to obtain lung slices to be used in motility experiments (see above) or (*ii*) infused i.v. with 5×10^6^ anti-EGFR CD8^+^ CAR T cells into the tail vein. After 4 days, mice were sacrificed, and lungs were collected. For flow cytometry, lungs were digested with DNase (Roche 4536282001) and Liberase (Roche 5401119001) solution at 37°C for 1h and then smashed using a 70µm Cell Strainer filter (Corning 352350). Collected cells were stained with Alexa647-anti-CD8 (Biolegend 344726) and PE-anti-EpCAM (Biolegend 324206) Abs for flow cytometry analysis (BD Accuri C6). For immunofluorescence, lungs were fixed in PLP solution (see above) for 24h at 4°C and, cut into 1-2mm^3^ pieces, embedded into 7% agarose (Sigma Aldrich A0701), and sliced using a vibratome (VT 1000S, Leica) to obtain 150µm-thick sections. Slices were processed using FITC-anti-CD3 (Biolegend 344804) and PE-anti-EPCAM (Biolegend 324206) Abs. To follow tumor growth, 3 days after i.v. injection of A549 cells, mice were injected i.v. with 5×10^5^ anti-EGFR CD8^+^ CAR T cells into the tail vein. Tumor growth in the lungs was followed by bioluminescence using PhotonIMAGER system (Biospacelab) and analysis was performed using M3Vision software (Biospacelab).

For the subcutaneous tumor model, 5×10^6^ human A549 cells were injected s.c. into the right flank of 2 months-old NSG nude mice. To evaluate CAR T cell infiltration, mice were injected i.v. after 5/6 weeks with 5×10^6^ anti-EGFR CD8^+^ CAR T cells and sacrificed 4 days after to perform the same analyses described for orthotopic tumors. Alternatively, 10 days after A549 cell injection, A549-derived tumor-bearing mice were i.v. inoculated with 5×10^6^ anti-EGFR CD8^+^ CAR T cells and tumor growth was followed through caliper measurement for additional 4 weeks. Then, mice were sacrificed, tumors were digested (like what described above for lungs), and collected cells were stained with PE-anti-EPCAM (Biolegend 324206) and Alexa488-anti-CD45 (Biolegend 304017) Abs for flow cytometry analysis (BD Accuri C6).

Subcutaneous tumors were also obtained through injection of human 5×10^6^ BxPC3 cells s.c. into the right flank of 2 months-old NSG nude mice (independently of sex). After 3/4 weeks, mice were sacrificed, and tumors were collected to obtain viable tumor slices used in the slice assay (see above).

### Western blot

Western blots were performed as previously described^78^. The following primary anti-human antibodies have been used: anti-actin (Cell Signaling 4970), anti-OGDH (Sigma Aldrich HPA019514), anti-Hsp90 (Cell Signaling 4877S), anti-PDHA1+A2 (Cell Signaling 2784S) and OxPhos Human WB Antibody Cocktail (Thermo Fisher 45-8199). Detection of protein signals was performed using ECL Prime Western Blotting Detection Reagent (Cytiva RPN2236) and Fusion FX instrument (Vilber Lourmat).

### Transendothelial migration assay

5µm-pore transwell inserts (Costar 3421) have been coated for 1h at 37°C with gelatin (Gibco S006100) and 10µg/ml bovine fibronectin protein (Thermo Fisher 33010018). Subsequently, 5×10^5^ HUVEC cells were added into the insert, let expanded for 24h and stimulated o.n. with 10ng/ml human TNFα (Thermo Fisher PHC3016). The day after, CAR T cells were stained with either Calcein Green or Cell Proliferation Dye eFluor670 (Thermo Fisher 65-0840-85) (same procedure used for motility assay, see above) and mixed at 1:1 ratio. Then, 2×10^5^ cells were added to the transwell upper chamber in a motility medium (BSA, no FBS). Transwell lower chamber was filled with RPMI complete medium (10% FBS). The transmigrated cells were collected and quantified by BD Accuri C6 cytometer.

### Statistical analysis & schemes in Figures

In the Figure legends, “n” indicates either the number of independent donors, the number of movies (no more than 3 from the same donor) or the number of mice used. Data are expressed as mean ± SEM or as boxplot. Comparisons between normal groups were done using two-tailed Student’s T-test (two groups) or One-way and Two-way ANOVA (multiple groups and repeated measurements, adjustments for pairwise comparisons were performed using Holm-Sidak method). Mann-Whitney Rank Sum Test or ANOVA on ranks were used if samples did not meet assumptions of normality. Significance is indicated in the Figures as follows: * = p < 0.05, ** = p < 0.01, *** = p < 0.001. Schemes shown in the figures have been created using BioRender.

## FUNDINGS

This work was partially supported by the European Union’s Horizon 2020 research and innovation programme under the Marie-Skłodowska Curie grant agreement No. 945298 “ParisRegionFP”, the French “Ligue Nationale contre le Cancer” (Equipes labellisées), GeFluc Ile-De-France (Entreprises Contre le Cancer), European Union’s Horizon Europe programme (HE-MSCA-PF 101062302), and Fondation ARC (PDF2 ARCPDF22020010001123). LS and MCAG were also supported by grant from Cochin Institut (PIC 2022). MF and DA were respectively supported by “Ligue Nationale contre le Cancer” and “Chinese Scientific Council” PhD fellowships.

## AUTHORS’ CONTRIBUTION

Conceptualization, Funding Acquisition & Writing - Original Draft: LS and ED. Investigation: LS, MC-AG, MF, DA, FP, and IR. Formal Analysis: LV. Resources: DD, ALM, MA, PI, DA and FP. Writing - Review & Editing: MC-AG, FP, PI. All authors approved the manuscript.

## CONFLICT OF INTEREST

Authors have no conflict of interest to declare.

## SUPPLEMENTARY MATERIALS

### Supplementary Figures

This PDF file contains Supplementary Figures 1-12.

### Supplementary Movies

This zip archive contains 58 movies of the 3D migration experiments shown throughout the manuscript (main figures, 2D z-stack reconstruction, 3 fps).

## Supporting information

Supplementary Informations

## Notes

### Competing Interest Statement

The authors have declared no competing interest.

## REFERENCES

1. Caruana, I., Simula, L., Locatelli, F. & Campello, S. T lymphocytes against solid malignancies: winning ways to defeat tumours. Cell Stress 2, 200–212 (2018).

2. Philip, M. & Schietinger, A. CD8+ T cell differentiation and dysfunction in cancer. Nat Rev Immunol 22, 209–223 (2022).

3. Raskov, H., Orhan, A., Christensen, J. P. & Gögenur, I. Cytotoxic CD8+ T cells in cancer and cancer immunotherapy. Br J Cancer 124, 359–367 (2021).

4. Joyce, J. A. & Fearon, D. T. T cell exclusion, immune privilege, and the tumor microenvironment. Science 348, 74–80 (2015).

5. Almagro, J., Messal, H. A., Elosegui-Artola, A., van Rheenen, J. & Behrens, A. Tissue architecture in tumor initiation and progression. Trends Cancer 8, 494–505 (2022).

6. Fu, T. et al. Spatial architecture of the immune microenvironment orchestrates tumor immunity and therapeutic response. J Hematol Oncol 14, 98 (2021).

7. Simsek, H. & Klotzsch, E. The solid tumor microenvironment—Breaking the barrier for T cells. BioEssays 44, 2100285 (2022).

8. Nicolas-Boluda, A. & Donnadieu, E. Obstacles to T cell migration in the tumor microenvironment. Comparative Immunology, Microbiology and Infectious Diseases 63, 22–30 (2019).

9. Salmon, H. et al. Matrix architecture defines the preferential localization and migration of T cells into the stroma of human lung tumors. Journal of Clinical Investigation 122, 899–910 (2012).

10. Peranzoni, E. et al. Macrophages impede CD8 T cells from reaching tumor cells and limit the efficacy of anti–PD-1 treatment. Proceedings of the National Academy of Sciences 115, E4041–E4050 (2018).

11. Simula, L., Ollivier, E., Icard, P. & Donnadieu, E. Immune Checkpoint Proteins, Metabolism and Adhesion Molecules: Overlooked Determinants of CAR T-Cell Migration? Cells 11, 1854 (2022).

12. Xiao, Z. et al. Disruption of desmoplastic stroma overcomes restrictions to T cell extravasation, immune exclusion and immunosuppression in solid tumors. 2023.04.13.536777 Preprint at 10.1101/2023.04.13.536777 (2023).

13. Simula, L. et al. Drp1 Controls Effective T Cell Immune-Surveillance by Regulating T Cell Migration, Proliferation, and cMyc-Dependent Metabolic Reprogramming. Cell Reports 25, 3059–3073 (2018).

14. Ledderose, C. et al. Purinergic P2X4 receptors and mitochondrial ATP production regulate T cell migration. The Journal of Clinical Investigation 128, 3583–3594 (2018).

15. Haas, R. et al. Lactate Regulates Metabolic and Pro-inflammatory Circuits in Control of T Cell Migration and Effector Functions. PLOS Biology 13, e1002202 (2015).

16. Campello, S. et al. Orchestration of lymphocyte chemotaxis by mitochondrial dynamics. J Exp Med 203, 2879–2886 (2006).

17. Chan, O., Burke, J. D., Gao, D. F. & Fish, E. N. The Chemokine CCL5 Regulates Glucose Uptake and AMP Kinase Signaling in Activated T Cells to Facilitate Chemotaxis. Journal of Biological Chemistry 287, 29406–29416 (2012).

18. Kishore, M. et al. Regulatory T Cell Migration Is Dependent on Glucokinase-Mediated Glycolysis. Immunity 47, 875–889.e10 (2017).

19. Le Bourgeois, T. et al. Targeting T Cell Metabolism for Improvement of Cancer Immunotherapy. Frontiers in Oncology 8, 237 (2018).

20. Stentz, F. B. & Kitabchi, A. E. Palmitic acid-induced activation of human T-lymphocytes and aortic endothelial cells with production of insulin receptors, reactive oxygen species, cytokines, and lipid peroxidation. Biochem Biophys Res Commun 346, 721–726 (2006).

21. Nava Lauson, C. B., et al. Linoleic acid potentiates CD8+ T cell metabolic fitness and antitumor immunity. Cell Metab 35, 633–650.e9 (2023).

22. Zhou, T. et al. Upregulation of SLAMF3 on human T cells is induced by palmitic acid through the STAT5-PI3K/Akt pathway and features the chronic inflammatory profiles of type 2 diabetes. Cell Death Dis 10, 559 (2019).

23. Angela, M. et al. Fatty acid metabolic reprogramming via mTOR-mediated inductions of PPARγ directs early activation of T cells. Nature Communications 7, 13683–13683 (2016).

24. Ioan-Facsinay, A. et al. Adipocyte-derived lipids modulate CD4+ T-cell function. Eur J Immunol 43, 1578–1587 (2013).

25. Lunt, S. Y. & Vander Heiden, M. G. Aerobic glycolysis: meeting the metabolic requirements of cell proliferation. Annu Rev Cell Dev Biol 27, 441–464 (2011).

26. Icard, P. & Simula, L. Metabolic oscillations during cell-cycle progression. Trends Endocrinol Metab 33, 447–450 (2022).

27. van der Windt, G. J. et al. Mitochondrial respiratory capacity is a critical regulator of CD8+ T cell memory development. Immunity 36, 68–78 (2012).

28. Bishop, E. L., Gudgeon, N. & Dimeloe, S. Control of T Cell Metabolism by Cytokines and Hormones. Front Immunol 12, 653605 (2021).

29. Rossignol, R. et al. Energy substrate modulates mitochondrial structure and oxidative capacity in cancer cells. Cancer Res 64, 985–993 (2004).

30. Marroquin, L. D., Hynes, J., Dykens, J. A., Jamieson, J. D. & Will, Y. Circumventing the Crabtree effect: replacing media glucose with galactose increases susceptibility of HepG2 cells to mitochondrial toxicants. Toxicol Sci 97, 539–547 (2007).

31. King, M. P. & Attardi, G. Human cells lacking mtDNA: repopulation with exogenous mitochondria by complementation. Science 246, 500–503 (1989).

32. Parri, M. & Chiarugi, P. Rac and Rho GTPases in cancer cell motility control. Cell Communication and Signaling 8, 23–23 (2010).

33. Hurd, T. R., DeGennaro, M. & Lehmann, R. Redox regulation of cell migration and adhesion. Trends Cell Biol 22, 107–115 (2012).

34. Xu, Q., Huff, L. P., Fujii, M. & Griendling, K. K. Redox regulation of the actin cytoskeleton and its role in the vascular system. Free Radic Biol Med 109, 84–107 (2017).

35. Aghajanian, A., Wittchen, E. S., Campbell, S. L. & Burridge, K. Direct activation of RhoA by reactive oxygen species requires a redox-sensitive motif. PLoS One 4, e8045 (2009).

36. Wolf, K., Müller, R., Borgmann, S., Bröcker, E.-B. & Friedl, P. Amoeboid shape change and contact guidance: T-lymphocyte crawling through fibrillar collagen is independent of matrix remodeling by MMPs and other proteases. Blood 102, 3262–3269 (2003).

37. Paňková, K., Rösel, D., Novotný, M. & Brábek, J. The molecular mechanisms of transition between mesenchymal and amoeboid invasiveness in tumor cells. Cell Mol Life Sci 67, 63–71 (2010).

38. Tantama, M., Martínez-François, J. R., Mongeon, R. & Yellen, G. Imaging energy status in live cells with a fluorescent biosensor of the intracellular ATP-to-ADP ratio. Nature Communications 4, 2550–2550 (2013).

39. Russo, E. et al. SPICE-Met: profiling and imaging energy metabolism at the single-cell level using a fluorescent reporter mouse. EMBO J 41, e111528 (2022).

40. He, S. et al. Characterization of the metabolic phenotype of rapamycin-treated CD8+ T cells with augmented ability to generate long-lasting memory cells. PLoS One 6, e20107 (2011).

41. Amiel, E. et al. Mechanistic target of rapamycin inhibition extends cellular lifespan in dendritic cells by preserving mitochondrial function. J Immunol 193, 2821–2830 (2014).

42. Araki, K. et al. mTOR regulates memory CD8 T-cell differentiation. Nature 460, 108–112 (2009).

43. Scholz, G. et al. Modulation of mTOR Signalling Triggers the Formation of Stem Cell-like Memory T Cells. EBioMedicine 4, 50–61 (2016).

44. Chang, C.-H. H. et al. Posttranscriptional Control of T Cell Effector Function by Aerobic Glycolysis. Cell 153, 1239–1251 (2013).

45. Chowdhury, P. S., Chamoto, K., Kumar, A. & Honjo, T. PPAR-Induced Fatty Acid Oxidation in T Cells Increases the Number of Tumor-Reactive CD8+ T Cells and Facilitates Anti-PD-1 Therapy. Cancer Immunol Res 6, 1375–1387 (2018).

46. Ma, E. H., Poffenberger, M. C., Wong, A. H.-T. & Jones, R. G. The role of AMPK in T cell metabolism and function. Curr Opin Immunol 46, 45–52 (2017).

47. Wang, Y. et al. NAD+ supplement potentiates tumor-killing function by rescuing defective TUB-mediated NAMPT transcription in tumor-infiltrated T cells. Cell Rep 36, 109516 (2021).

48. Hong, Y. et al. ST3GAL1 and βII-spectrin pathways control CAR T cell migration to target tumors. Nat Immunol 24, 1007–1019 (2023).

49. Forster, J. C., Harriss-Phillips, W. M., Douglass, M. J. & Bezak, E. A review of the development of tumor vasculature and its effects on the tumor microenvironment. Hypoxia (Auckl) 5, 21–32 (2017).

50. Sommermeyer, D. et al. Chimeric antigen receptor-modified T cells derived from defined CD8+ and CD4+ subsets confer superior antitumor reactivity in vivo. Leukemia 30, 492–500 (2016).

51. Xu, Y. et al. Closely related T-memory stem cells correlate with in vivo expansion of CAR.CD19-T cells and are preserved by IL-7 and IL-15. Blood 123, 3750–3759 (2014).

52. Louis, C. U. et al. Antitumor activity and long-term fate of chimeric antigen receptor-positive T cells in patients with neuroblastoma. Blood 118, 6050–6056 (2011).

53. O’Sullivan, D. et al. Fever supports CD8+ effector T cell responses by promoting mitochondrial translation. Proceedings of the National Academy of Sciences 118, e2023752118 (2021).

54. Shiraishi, T. et al. Glycolysis is the primary bioenergetic pathway for cell motility and cytoskeletal remodeling in human prostate and breast cancer cells. Oncotarget 6, 130–143 (2015).

55. Zanotelli, M. R., Zhang, J. & Reinhart-King, C. A. Mechanoresponsive metabolism in cancer cell migration and metastasis. Cell Metab 33, 1307–1321 (2021).

56. Porporato, P. E. et al. A Mitochondrial Switch Promotes Tumor Metastasis. Cell Reports 8, 754–766 (2014).

57. Amitrano, A. M. et al. Optical Control of CD8+ T Cell Metabolism and Effector Functions. Front Immunol 12, 666231 (2021).

58. Sena, L. A. et al. Mitochondria Are Required for Antigen-Specific T Cell Activation through Reactive Oxygen Species Signaling. Immunity 38, 225–236 (2013).

59. Franchina, D. G., Dostert, C., Brenner, D., Lu, D. B. & Brenner, D. Reactive Oxygen Species: Involvement in T Cell Signaling and Metabolism. Trends in Immunology 39, 489–502 (2018).

60. Dumauthioz, N. et al. Enforced PGC-1α expression promotes CD8 T cell fitness, memory formation and antitumor immunity. Cell Mol Immunol 18, 1761–1771 (2021).

61. Lontos, K. et al. Metabolic reprogramming via an engineered PGC-1α improves human chimeric antigen receptor T-cell therapy against solid tumors. J Immunother Cancer 11, e006522 (2023).

62. Crompton, J. G. et al. Akt inhibition enhances expansion of potent tumor-specific lymphocytes with memory cell characteristics. Cancer Res 75, 296–305 (2015).

63. Zhang, Y. et al. Enhancing CD8+ T Cell Fatty Acid Catabolism within a Metabolically Challenging Tumor Microenvironment Increases the Efficacy of Melanoma Immunotherapy. Cancer Cell 32, 377–391.e9 (2017).

64. Rostamian, H., Khakpoor-Koosheh, M., Fallah-Mehrjardi, K., Mirzaei, H. R. & Brown, C. E. Mitochondria as Playmakers of CAR T-cell Fate and Longevity. Cancer Immunol Res 9, 856–861 (2021).

65. Pellegrino, M. et al. Manipulating the Metabolism to Improve the Efficacy of CAR T-Cell Immunotherapy. Cells 10, 14 (2020).

66. Peng, J.-J., Wang, L., Li, Z., Ku, C.-L. & Ho, P.-C. Metabolic challenges and interventions in CAR T cell therapy. Sci Immunol 8, eabq3016 (2023).

67. Al Tameemi, W., Dale, T. P., Al-Jumaily, R. M. K. & Forsyth, N. R. Hypoxia-Modified Cancer Cell Metabolism. Front Cell Dev Biol 7, 4 (2019).

68. Kierans, S. J. & Taylor, C. T. Regulation of glycolysis by the hypoxia-inducible factor (HIF): implications for cellular physiology. J Physiol 599, 23–37 (2021).

69. Scharping, N. E. et al. Mitochondrial stress induced by continuous stimulation under hypoxia rapidly drives T cell exhaustion. Nat Immunol 22, 205–215 (2021).

70. Rytelewski, M. et al. Merger of dynamic two-photon and phosphorescence lifetime microscopy reveals dependence of lymphocyte motility on oxygen in solid and hematological tumors. J Immunother Cancer 7, 78 (2019).

71. Khacho, M. et al. Acidosis overrides oxygen deprivation to maintain mitochondrial function and cell survival. Nat Commun 5, 3550 (2014).

72. Lim, A. R., Rathmell, W. K. & Rathmell, J. C. The tumor microenvironment as a metabolic barrier to effector T cells and immunotherapy. Elife 9, e55185 (2020).

73. Elia, I. et al. Tumor cells dictate anti-tumor immune responses by altering pyruvate utilization and succinate signaling in CD8+ T cells. Cell Metab 34, 1137–1150.e6 (2022).

74. Quinn, W. J. et al. Lactate Limits T Cell Proliferation via the NAD(H) Redox State. Cell Rep 33, 108500 (2020).

75. Watson, M. J. et al. Metabolic support of tumour-infiltrating regulatory T cells by lactic acid. Nature 591, 645–651 (2021).

76. Peranzoni, E. et al. Ex Vivo Imaging of Resident CD8 T Lymphocytes in Human Lung Tumor Slices Using Confocal Microscopy. Journal of Visualized Experiments e57709–e57709 (2017) doi:10.3791/55709.

77. Ershov, D. et al. TrackMate 7: integrating state-of-the-art segmentation algorithms into tracking pipelines. Nat Methods 19, 829–832 (2022).

78. Simula, L. et al. PD-1-induced T cell exhaustion is controlled by a Drp1-dependent mechanism. Mol Oncol 16, 188–205 (2022).

