## Supplementary Informations for "Mitochondrial metabolism sustains CD8^+^ T cell migration for an efficient infiltration into solid tumors"

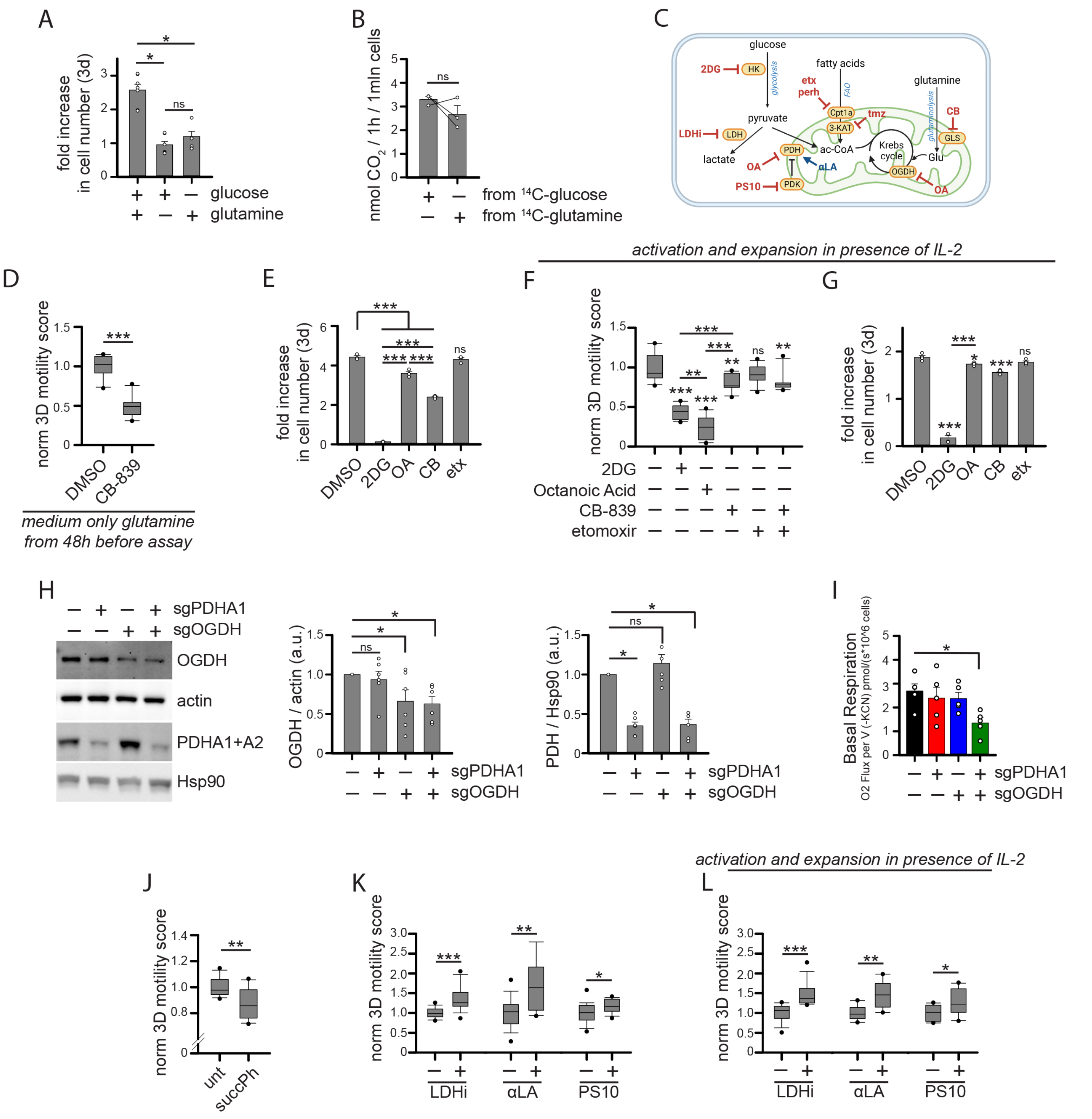


**Supplementary Figure 1. Further analyses about the metabolic regulation of T cell 3D motility. Related to Figure 1.**
(**A**) Fold increase in cell number of activated CD8+ T cells cultured for 3d in absence of glucose or glutamine (n=7).
(**B**) $CO_2$ production from radio-labelled glucose or glutamine in activated CD8+ T cells (n=3).
(**C**) Scheme of drugs used to inhibit or activate metabolic enzymes and regulatory proteins.
(**D**) Normalized 3D motility Score in presence of CB-839 of activated CD8+ T cells cultured since 48h before assay in medium containing glutamine and no glucose (n=10).
(**E**) Fold increase in cell number of activated CD8+ T cells cultured for 3d in presence of the indicated drugs (n=3).
(**F-G**) Normalized 3D motility Score (F, n=11) and fold increase in cell number after 3d of culture (G, n=4) in presence of the indicated drugs for activated CD8+ T cells cultured in presence of IL-2.
(**H**) WB analysis of protein expression in activated CD8+ T cells CRISPR/Cas9-edited for the indicated genes (n=6).
(**I**) Basal respiration in activated CD8+ T cells CRISPR/Cas9-edited for the indicated genes (n=5).
(**J**) Normalized 3D motility Score of activated CD8+ T cells in presence of OGDH inhibitor succinyl-phosphonate (succPh) (n=12).
(**K-L**) Normalized 3D motility Score of activated CD8+ T cells in presence of the indicated drugs. Since initial activation, cells were cultured with IL7+IL15 (L, LDHi n=18, αLA n=15, PS10 n=21) or IL2 (M, LDHi & PS10 n=13, αLA n=9).
Data are shown as mean ± SEM or box plot. Significance is indicated as follows: ns = not significant, * = p<0.05, ** = p<0.01, *** = p<0.001.

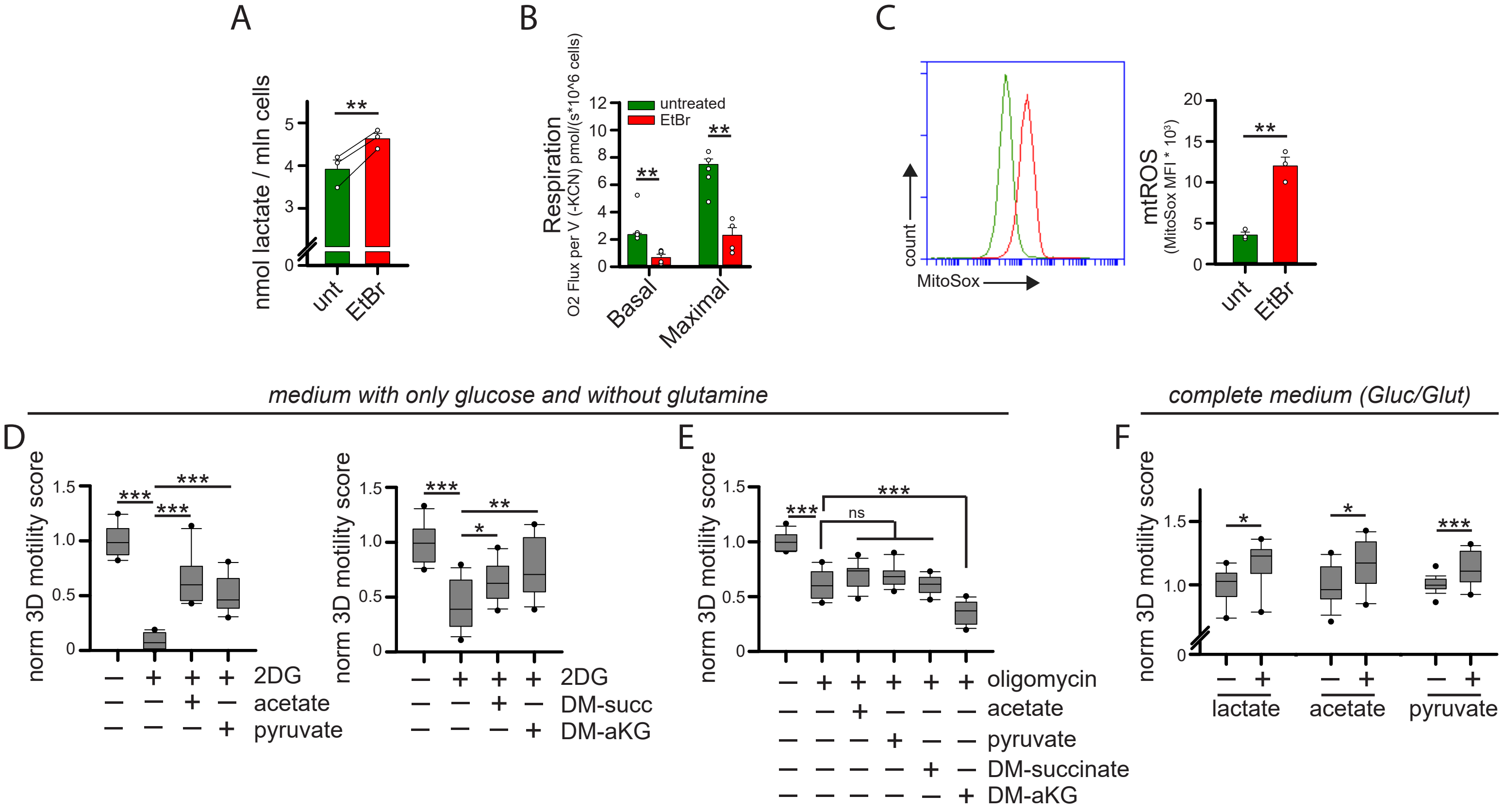


**Supplementary Figure 2. Dissecting glycolysis/OXPHOS importance to sustain T cell 3D motility. Related to Figure 2.**
(**A-C**) Activated CD8⁺ T cells were cultured in presence of ethidium bromide (EtBr) for at least 14d. Then, the amount of lactate (A, n=3), the basal and maximal respiration (B, n=4) and the amount of mtROS (MitoSox/FSC ratio; C, n=3) have been evaluated.
(**D-E**) Normalized 3D motility Score of activated CD8⁺ T cells in presence of the indicated drugs (2DG in D, n=10 left graph, n=12 right graph; oligomycin in E, n=12) and nutrients (dimethyl-succinate: DM-succ; dimethyl-alpha-ketoglutarate: DM-aKG) in a motility medium containing only glucose and no glutamine.Please note that data from rows 1-2 of figure S2E were also used in figure 2I for lactate since performed in a single whole experiment (data were splitted to improve quality of the presentation).
(**F**) Normalized 3D motility Score of activated CD8⁺ T cells in presence of the indicated nutrients in a complete motility medium (glucose + glutamine) (lactate n=9, acetate n=12, pyruvate n=19).
Data are shown as mean ± SEM or boxplot. Significance is indicated as follows: ns = not significant, * = p<0.05, ** = p<0.01, *** = p<0.001.

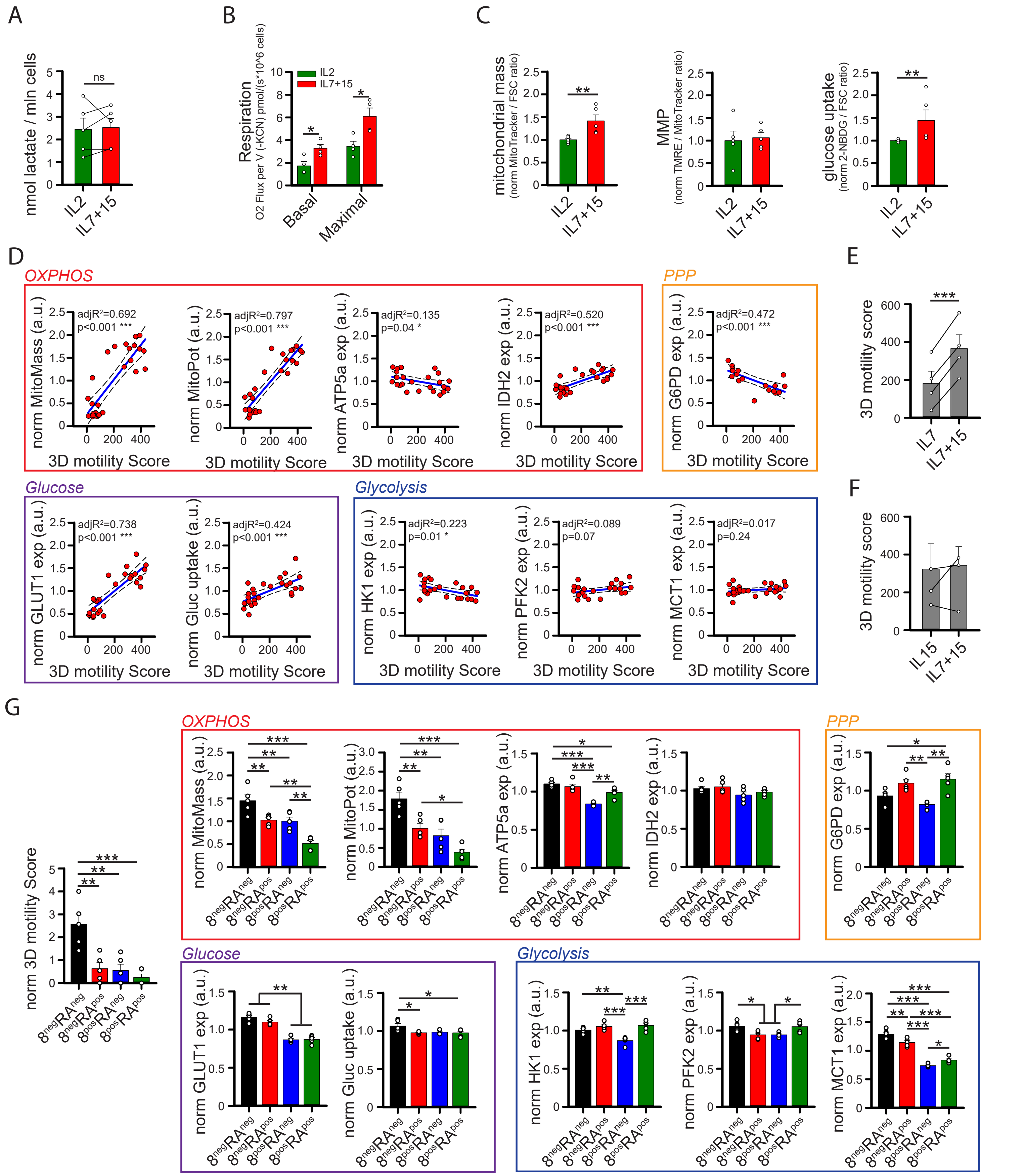


**Supplementary Figure 3. Further correlations between T cell motility and mitochondrial metabolism. Related to Figure 3.**
(**A-C**) CD8⁺ T cells have been activated and cultured in presence of IL2 or IL7+IL15 (n=7). Then, the amount of lactate (A, n=5), the basal and maximal respiration (B, n=4), the amount of mitochondria (MitoTrackerGreen/FSC ratio C, n=5), the mitochondrial membrane potential (MMP, TMRE/MitoTrackerGreen ratio; C, n=5) and the amount of mtROS (MitoSox/FSC ratio; C, n=5) have been evaluated.
(**D**) CD8⁺ T cells were activated and cultured in presence of varying doses of IL7+15, IL2, and a-CD3/28 Abs. After 1 week, 3D motility and the level (MFI, flow cytometry) of indicated proteins or parameters (MitoMass: MitoTrackerGreen/FSC ratio; MitoPot: TMRE/MitoTrackerGreen ratio; Gluc uptake: 2-NBDG/FSC ratio) were evaluated. The graphs show the correlation between the normalized 3D motility Score and the indicated parameters (n=18 for G6PD, n=25 for ATP5a, GLUT1, IDH2 and HK1, n=26 for others).
(**E-F**) CD8⁺ T cells were activated and cultured in presence of IL7, IL15 or IL7+15. Then, motility onto BxPC3-derived tumor slice was evaluated as in Figure 3C (E, n=4; F, n=3).
(**G**) 3D motility and the level (MFI) of indicated proteins or parameters (as in D) were evaluated in freshly isolated hPBT cells not stimulated and stained with CD8 and CD45RA Abs to distinghuish the indicated populations (n=5).
Data are shown as mean ± SEM or box plot. Significance is indicated as follows: **= p<0.05, ** = p<0.01, *** = p<0.001.

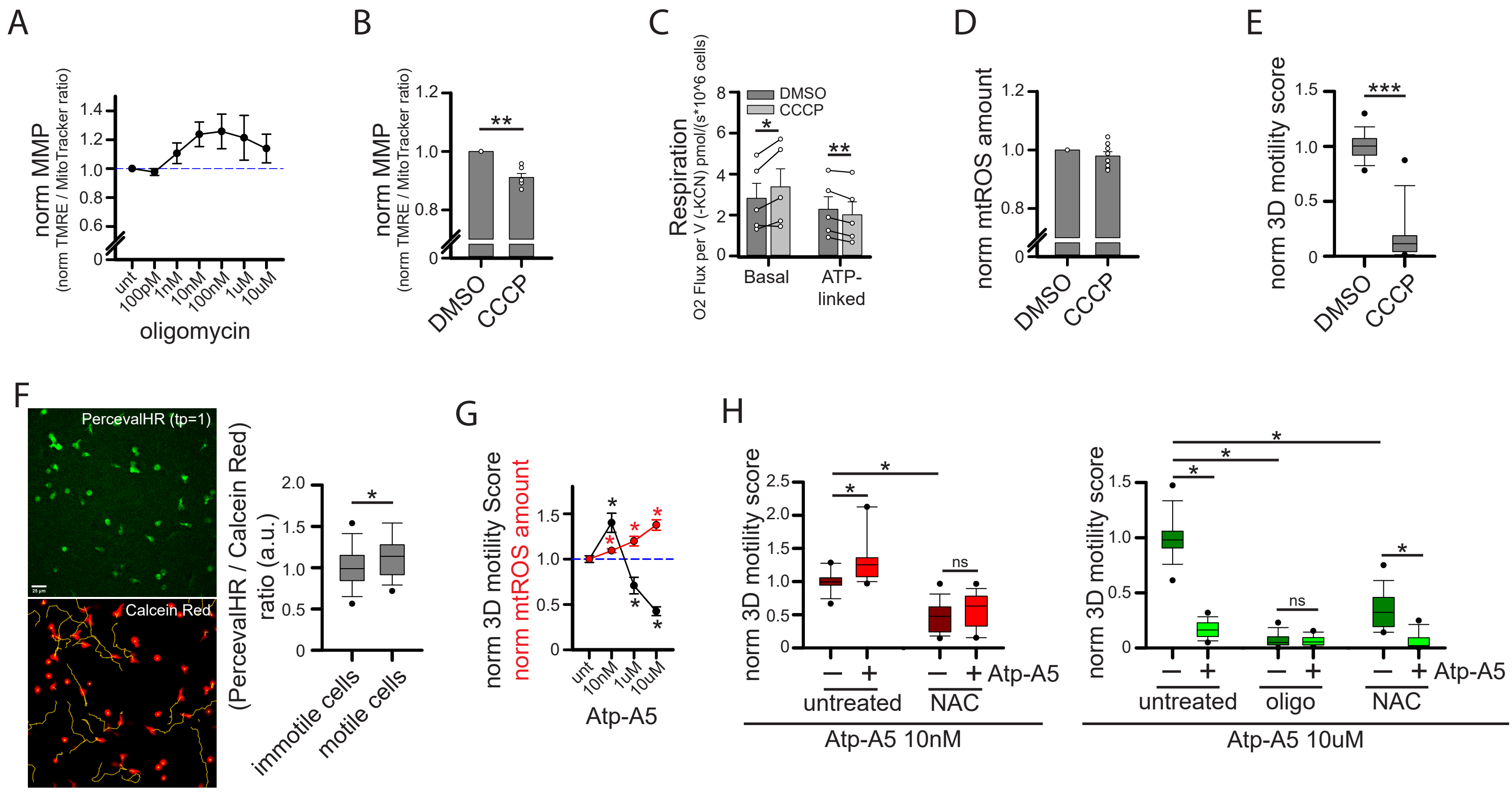


**Supplementary Figure 4. Further analyses about the role of ATP and mtROS to sustain 3D motility. Related to Figure 4.**
(**A**) Normalized mitochondrial membrane potential (MMP, TMRE/MitoTrackerGreen ratio) in activated CD8$^+$ T cells cultured for 1h in presence of the indicated doses of oligomycin (n=5).
(**B-E**) Activated CD8$^+$ T cells were cultured in presence of CCCP for 1h. Then, normalized mitochondrial membrane potential (MMP, TMRE/MitoTrackerGreen ratio; B, n=6), basal and ATP-linked respiration (C n=5), mtROS amount (Mitosox/FSC ratio; D, n=8) and normalized 3D motility Score (E, n=15) were evaluated.
(**F**) Activated CD8$^+$ T cells have been infected with lentiviral particles to express the ATP sensor PercevalHR. After 1w, cells were stained with Calcein Red and 3D motility assessed in collagen gel. PercevalHR MFI was calculated for each cell at first timepoint (tp=1) and then motility evaluated by TrackMate. Cells were distinghuished into immotile and motile cells according to their displacement and the relative PercevalHR/CalceinRed ratio calculated as a proxy to infer ATP amount (n=62 immotile cells and n=45 motile cells).
(**G**) Normalized 3D motility Score (n=15) and mtROS amount (Mitosox/FSC ratio; n=6) were evaluated in activated CD8$^+$ T cells in presence of the indicated doses of the ETC-II complex inhibitor Atpenin-A5 (Atp-A5).
(**H**) Normalized 3D motility Score of activated CD8$^+$ T cells evaluated in presence of the indicated doses of Atpenin-A5 (Atp-A5), N-acetyl-cysteine (NAC) and oligomycin (oligo) (n=18).
Data are shown as mean ± SEM or box plot. Significance is indicated as follows: ns = not significant, * = p<0.05, ** = p<0.01, *** = p<0.001.

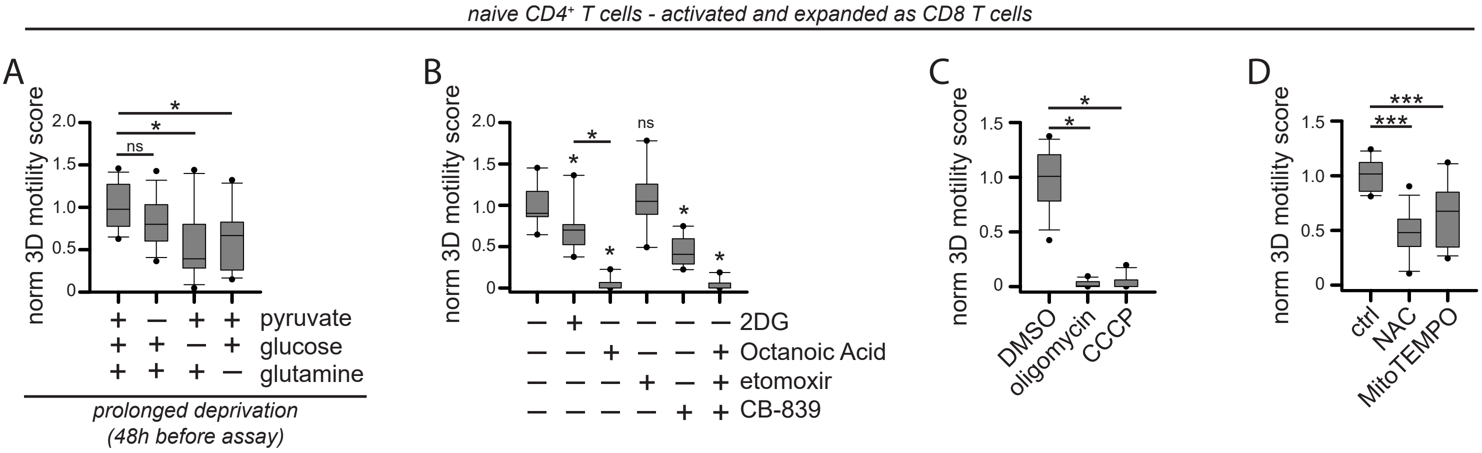


**Supplementary Figure 5. Mitochondrial metabolism support CD4⁺ T cell 3D motility.**
CD4⁺ T cells have been activated and expanded as CD8⁺ T cells (IL7+15).
(**A**) Normalized 3D motility Score of activated CD4⁺ T cells cultured from 48h before assay in medium containing no glucose, no pyruvate or no glutamine (n=12).
(**B-D**) Normalized 3D motility Score of activated CD4⁺ T cells in presence of the indicated drugs (B, n=9; C, n=12; D, n=12).
Data are shown as box plot. Significance is indicated as follows: ns = not significant, * = p<0.05, ** = p<0.01, *** = p<0.001.

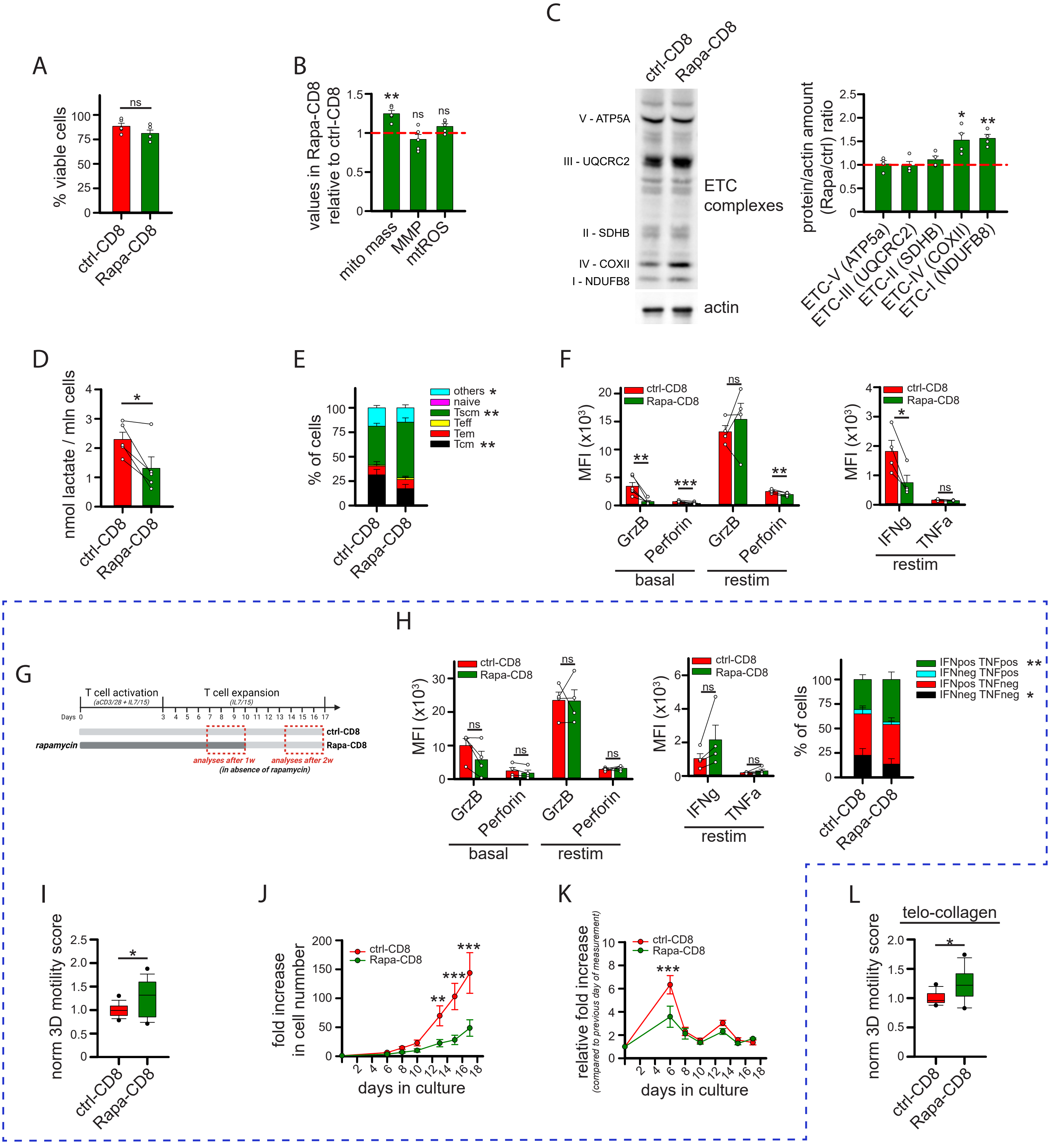


**Supplementary Figure 6. Further analyses in rapamycin-treated CD8⁺ T cells. Related to Figure 5.**
(**A-F**) Activated CD8⁺ T cells have been cultured in ctrl condition (ctrl-CD8) or in presence of rapamycin (Rapa-CD8). After 1w, rapamycin was removed the day of the analysis and the following parameters were measured: percentage of viable cells (A, n=5), the amount of mitochondria (mito mass: MitoTrackerGreen/FSC ratio in Rapa-CD8 relative to ctrl-CD8; B, n=5), the mitochondrial membrane potential (MMP: TMRE/MitoTrackerGreen ratio in Rapa-CD8 relative to ctrl-CD8; B, n=5), the amount of mtROS (MitoSox/FSC ratio in Rapa-CD8 relative to ctrl-CD8; B, n=5), the expression of the indicated proteins in Rapa-CD8 relative to ctrl-CD8 (WB, C, n=4), lactate production (D, n=5), the relative proportion of indicated subsets (flow cytometry, E, n=6), and the level (MFI) of the indicated protein in resting (basal) or restimulated (restim, 5h with human TransAct + BfdA) cells (F, n=6 basal, n=4 restim).
(**G-K**) Activated CD8⁺ T cells have been cultured in control condition (ctrl-CD8) or in presence of rapamycin (Rapa-CD8) for 10d. Then, rapamycin was removed and cells were expanded in control conditions for additional 7d (G, scheme). The following analyses was performed at 14-17d: the level (MFI) of the indicated protein in resting cells (basal) or in cells restimulated (restim) 5h with human TransAct + BfdA (H, for IFNg and TNFa the percentage of cells producing these cytokines are indicated in the graph on the right; n=4), the normalized 3D motility Score (I, n=27), the fold increase in cell number (J, n=5), and the relative fold increase in cell number calculated as the fold increase at day "x" compared to previous day of measurement (K, n=5).
(**L**) Normalized 3D motility Score of activated CD8⁺ T cells in 3mg/ml telocollagen gel (n=12).
Data are shown as mean ± SEM or box plot. Significance is indicated as follows: ns = not significant, * = p<0.05, ** = p<0.01, *** = p<0.001.

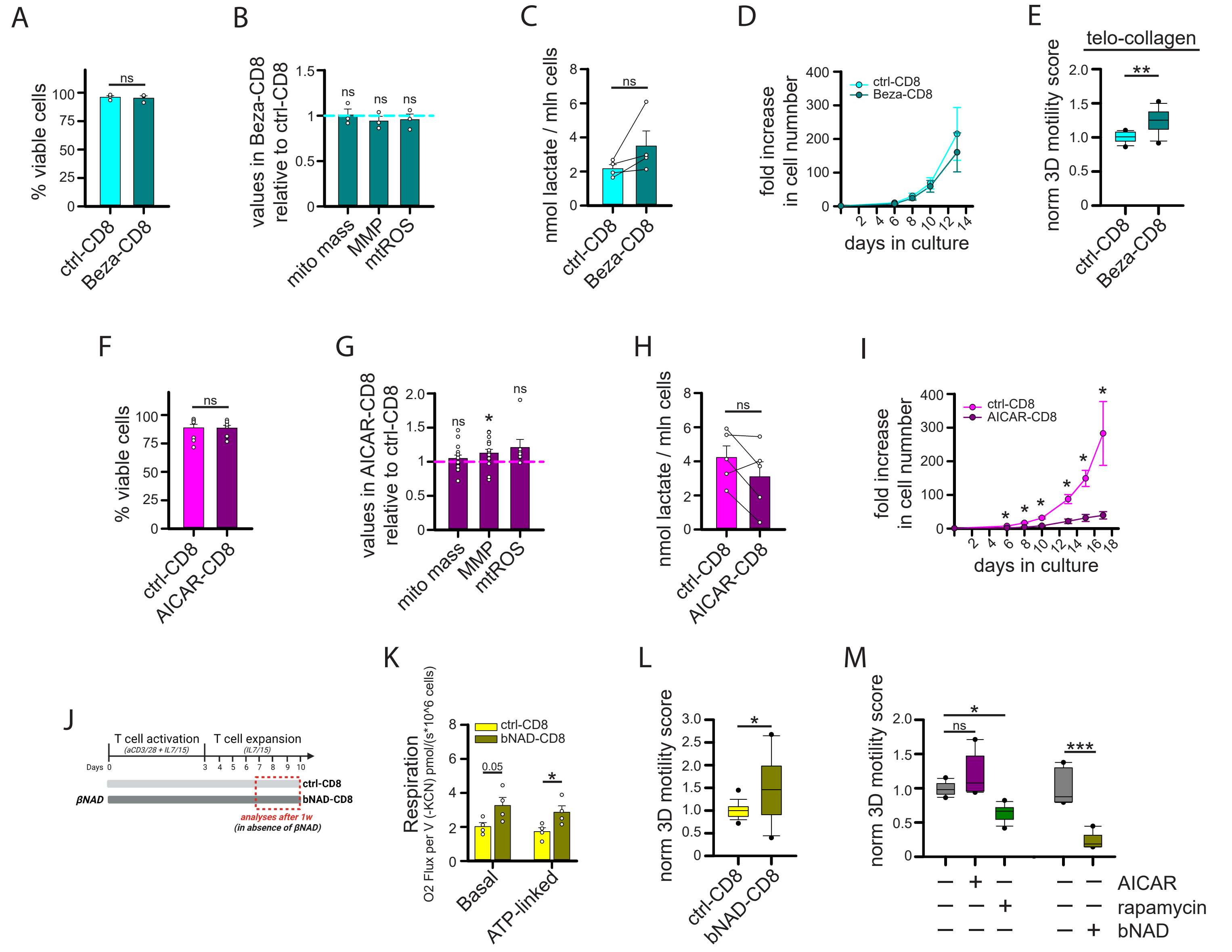


**Supplementary Figure 7. Further analyses in bezafibrate- and AICAR-treated CD8⁺ T cells. Related to Figure 5.**
(**A-E**) Activated CD8⁺ T cells have been cultured in ctrl condition (ctrl-CD8) or in presence of bezafirbate (Beza-CD8). After 1w, bezafibrate was removed the day of the analysis and the following parameters were measured: percentage of viable cells (A, n=3), the amount of mitochondria (mito mass: MitoTrackerGreen/FSC ratio in Beza-CD8 relative to ctrl-CD8; B, n=3), the mitochondrial membrane potential (MMP: TMRE/MitoTrackerGreen ratio in Beza-CD8 relative to ctrl-CD8; B, n=3), the amount of mtROS (MitoSox/FSC ratio in Beza-CD8 relative to ctrl-CD8; B, n=3), and lactate production (C, n=4).
(**D**) Fold increase in cell number for ctrl-CD8 and Beza-CD8 T cells (obtained as in A). Bezafibrate was removed at 10d (n=4).
(**E**) Normalized 3D motility Score of ctrl-CD8 and Beza-CD8 T cells (obtained as in A) in 3mg/ml telocollagen gel (n=12).
(**F-H**) Activated CD8⁺ T cells have been cultured in ctrl condition (ctrl-CD8) or in presence of AICAR (AICAR-CD8). After 3d, AICAR was removed and the cells expanded in control condition. The following parameters were measured starting from 7d: percentage of viable cells (F, n=9), the amount of mitochondria (mito mass: MitoTrackerGreen/FSC ratio in AICAR-CD8 relative to ctrl-CD8; G, n=15), the mitochondrial membrane potential (MMP: TMRE/MitoTrackerGreen ratio in AICAR-CD8 relative to ctrl-CD8; G, n=15), the amount of mtROS (MitoSox/FSC ratio in AICAR-CD8 relative to ctrl-CD8; G, n=7), and lactate production (H, n=5).
(**I**) Fold increase in cell number for ctrl-CD8 and AICAR-CD8 T cells (obtained as in F). AICAR was removed at 3d (n=4).
(**J-L**) CD8⁺ T cells have been cultured in ctrl condition (ctrl-CD8) or in presence of 1mM βNAD (bNAD-CD8). After 1w, βNAD was removed the day of the analysis (J, scheme) and the following parameters were measured: basal and ATP-linked respiration (K, n=4) and normalized 3D motility Score (L, n=18).
(**M**) Normalized 3D motility Score of control activated CD8⁺ T cells in presence of the indicated drugs during the assay (AICAR & Rapa, n=12; bNAD, n=6)
Data are shown as mean ± SEM or box plot. Significance is indicated as follows: ns = not significant, * = $p<0.05$, ** = $p<0.01$, *** = $p<0.001$.

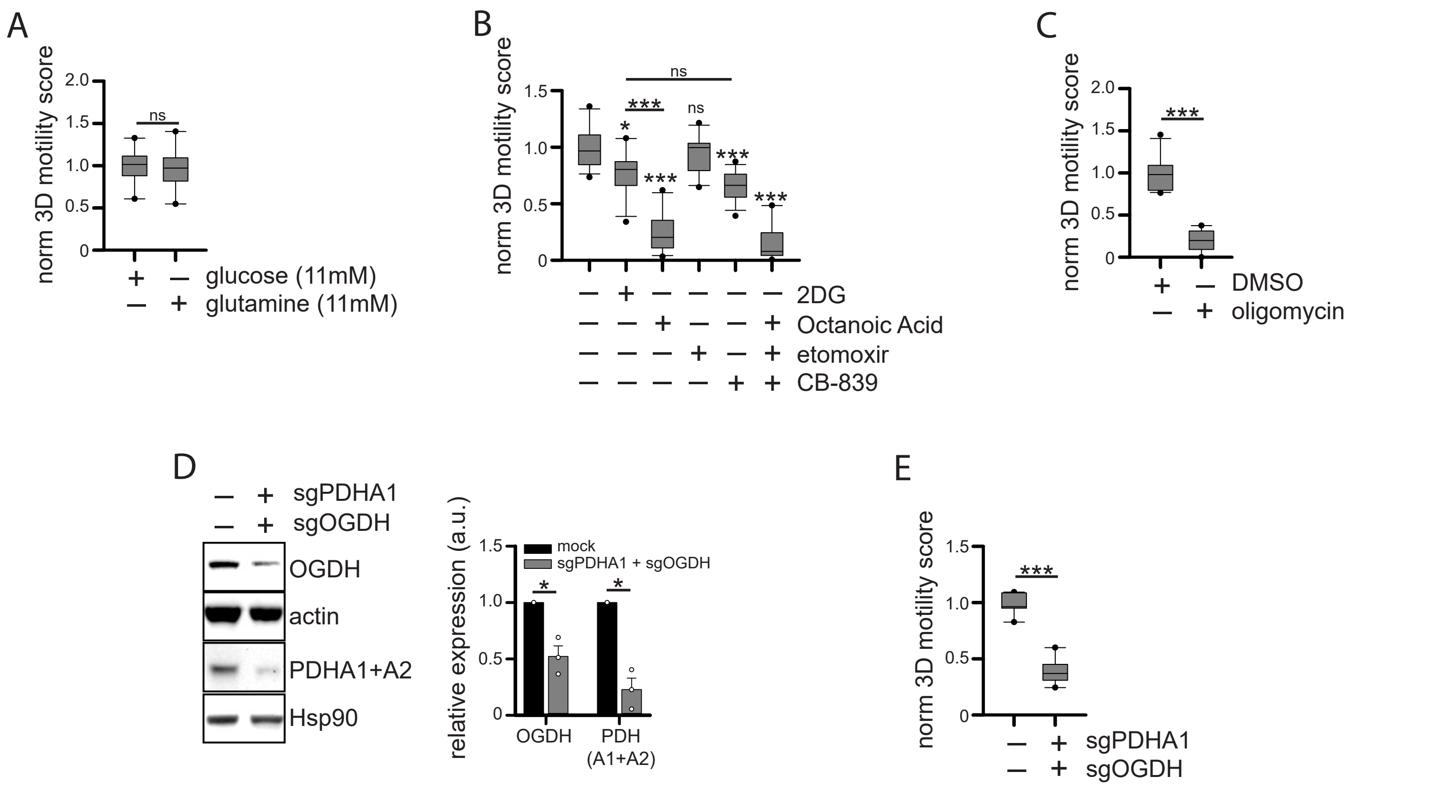


**Supplementary Figure 8. Mitochondrial metabolism supports anti-EGFR CD8+ CAR T cell 3D motility.**
(**A-C**) Activated CD8+ T cells have been infected with lentiviral particles to generate CD8+ CAR T cells recognizing the EGFR antigen (anti-EGFR CD8+ CAR T cells). After 1w, normalized 3D motility Score was calculated in motility medium containing no glucose or no glutamine (A, n=8) or in presence of the indicated drugs (B, n=12; C, n=12).
(**D-E**) Anti-EGFR CD8+ CAR T cells (obtained as in A) were CRISPR/Cas9-edited CRISPR/Cas9-edited for the indicated genes. After 1w, the levels of the indicated protein (WB analysis; D, n=3) and the normalized 3D motility Score (E, n=9) were evaluated. Data are shown as mean ± SEM or box plot. Significance is indicated as follows: ns = not significant, * = p<0.05, ** = p<0.01, *** = p<0.001.

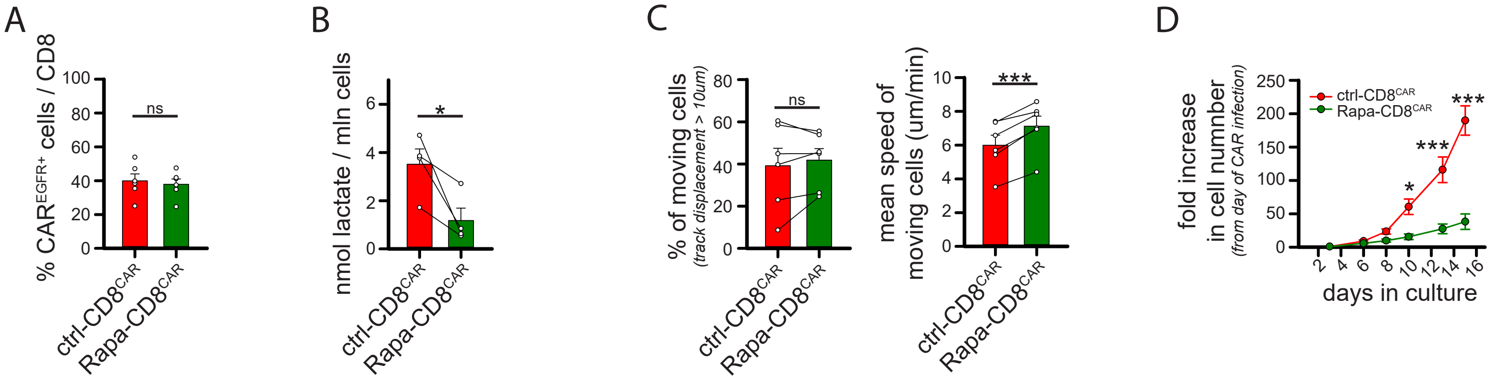


**Supplementary Figure 9. Mitochondrial metabolism supports anti-EGFR $CD8^+$ CAR T cell 3D motility. Related to Figure 6.** (**A-C**) Activated $CD8^+$ T cells have been infected with lentiviral particles to generate $CD8^+$ CAR T cells recognizing the EGFR antigen (anti-EGFR $CD8^+$ CAR T cells) and cultured in control conditions (ctrl-$CD8^{CAR}$) or in presence of rapamycin (Rapa-$CD8^{CAR}$). After 1w, rapamycin was removed the day of the analysis and the following parameters were measured: percentage of cells expressing the anti-EGFR CAR molecule at the surface (A, n=6), lactate amount (B, n=4), and the two parameters composing the 3D motiity Score: percentage of moving cells and mean speed of moving cells calculated as mean values for each donor (C, n=6). (**D**) Fold increase in cell number for ctrl-$CD8^{CAR}$ and Rapa-$CD8^{CAR}$ T cells (obtained as in A). Rapa was removed at 10d (n=4). Data are shown as mean ± SEM or box plot. Significance is indicated as follows: ns = not significant, * = $p<0.05$, ** = $p<0.01$, *** = $p<0.001$.

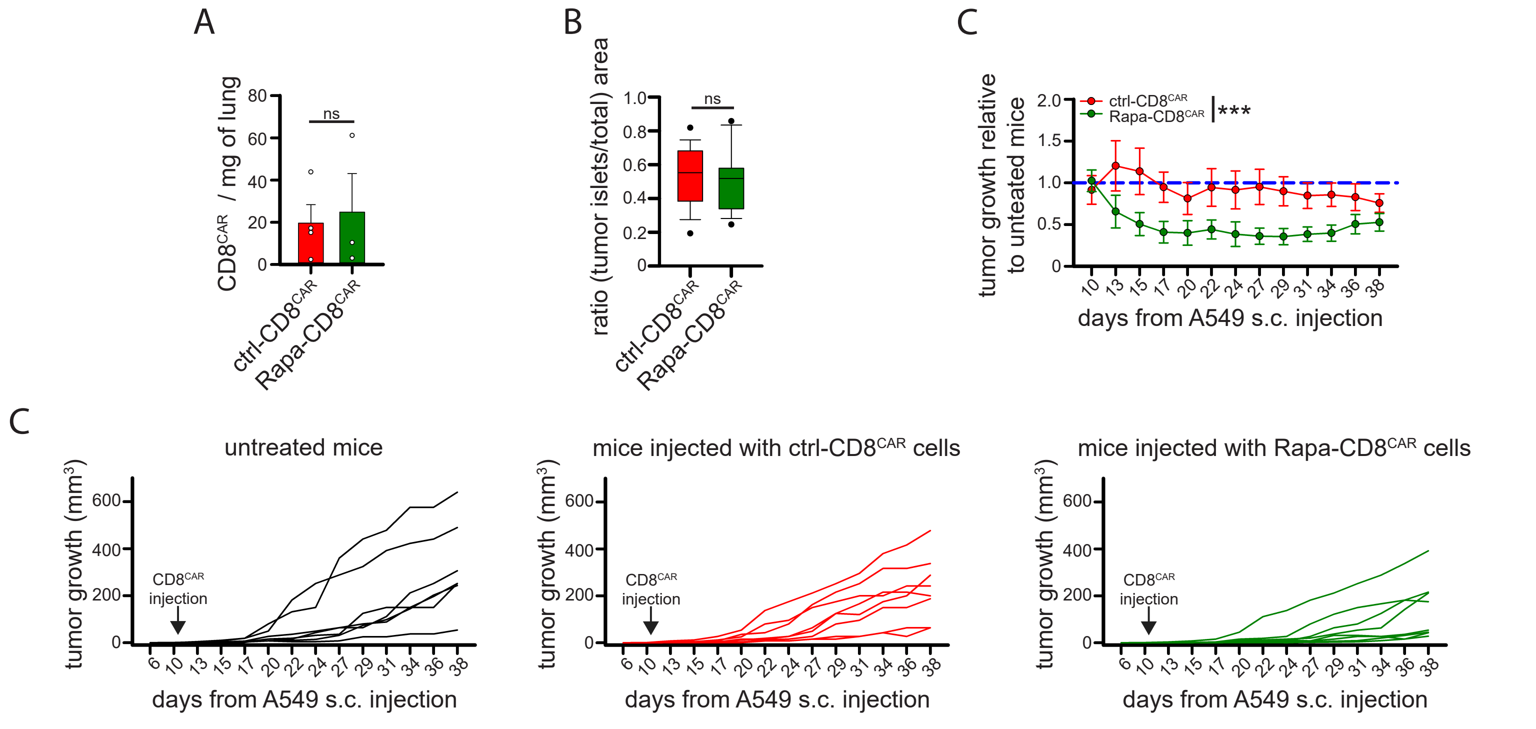


**Supplementary Figure 10. Further *in vivo* analyses of the efficacy of Rapa-CD8$^{CAR}$ T cells. Related to Figure 7.**

(**A**) Number of anti-EGFR CD8$^{+}$ CAR T cells (per mg of lung) recoverd (flow cytometry) from lung of NSG mice i.v. injected with A549 cells 3-weeks before and i.v. injected with ctrl-CD8$^{CAR}$ or Rapa-CD8$^{CAR}$ cells 4d before (see scheme in figure 7A) (ctrl-CD8$^{CAR}$, n=4; Rapa-CD8$^{CAR}$, n=3).

(**B**) Same experiment as in Figure 7D. NSG mice were injected s.c. with with A549 cells and then, 5/6w after, injected i.v. with ctrl-CD8$^{CAR}$ or Rapa-CD8$^{CAR}$ cells. After 4d, mice were sacrificed, and tumors isolated, fixed and cut through vibratome. Here, it is reported the analysis of the relative area of each section occupied by tumor cells (EpCAM$^{+}$) obtained from immunofluorescence images as shown in Figure 7D.

(**C-D**) Same experiment as in Figure 7E. Tumor growth was measured in NSG mice s.c. injected with A549 cells at indicated days since A549 cell injection. Ctrl-CD8$^{CAR}$ or Rapa-CD8$^{CAR}$ cells was injected i.v. at 10d. In C (n=8), it is reported the size of tumors growing in NSG mice injected with ctrl-CD8$^{CAR}$ or Rapa-CD8$^{CAR}$ cells relative to uninjected tumor-bearing NSG mice in the same cage (mean tumor size of untreated mice group set to 1 for each cage). In D (untreated, n=7; ctrl-CD8$^{CAR,}$ n=8; Rapa-CD8$^{CAR}$, n=8), the single growth curves of tumors in untreated NSG mice (left) or NSG mice injected with ctrl-CD8$^{CAR}$ (middle) or Rapa-CD8$^{CAR}$ cells (right) are reported.

Data are shown as mean ± SEM. Significance is indicated as follows: ns = not significant, * = p<0.05, ** = p<0.01, *** = p<0.001.

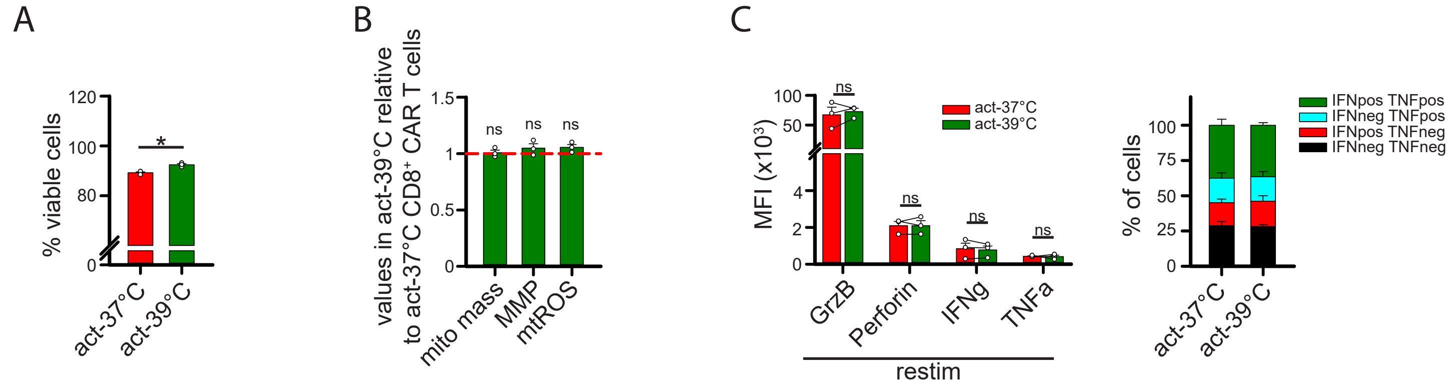


**Supplementary Figure 11. Further analyses in CD8$^+$ CAR T cells activated at 39°C. Related to Figure 8.**
(**A-C**) CD8$^+$ T cells have been activated at 37°C (act-37°C) or at 39°C (act-39°C) for 3d. Then, cells have been put at 37°C and infected with lentiviral particles to generate CD8$^+$ CAR T cells recognizing the EGFR antigen (anti-EGFR CD8$^+$ CAR T cells) and subsequently expanded at 37°C. After 1w, the following parameters were measured: percentage of viable cells (A, n=3), the amount of mitochondria (mito mass: MitoTrackerGreen/FSC ratio in act-39°C relative to act-37°C; B, n=3), the mitochondrial membrane potential (MMP: TMRE/MitoTrackerGreen ratio in act-39°C relative to act-37°C; B, n=3), the amount of mtROS (MitoSox/FSC ratio in act-39°C relative to act-37°C; B, n=3), and the level (MFI) of the indicated protein in restimulated (restim, 5h with human TransAct + BfdA) cells (for IFNg and TNFa the percentage of cells producing these cytokines are indicated in the graph on the right) (C, n=3).
Data are shown as mean ± SEM or box plot. Significance is indicated as follows: ns = not significant, * = p<0.05, ** = p<0.01, *** = p<0.001.

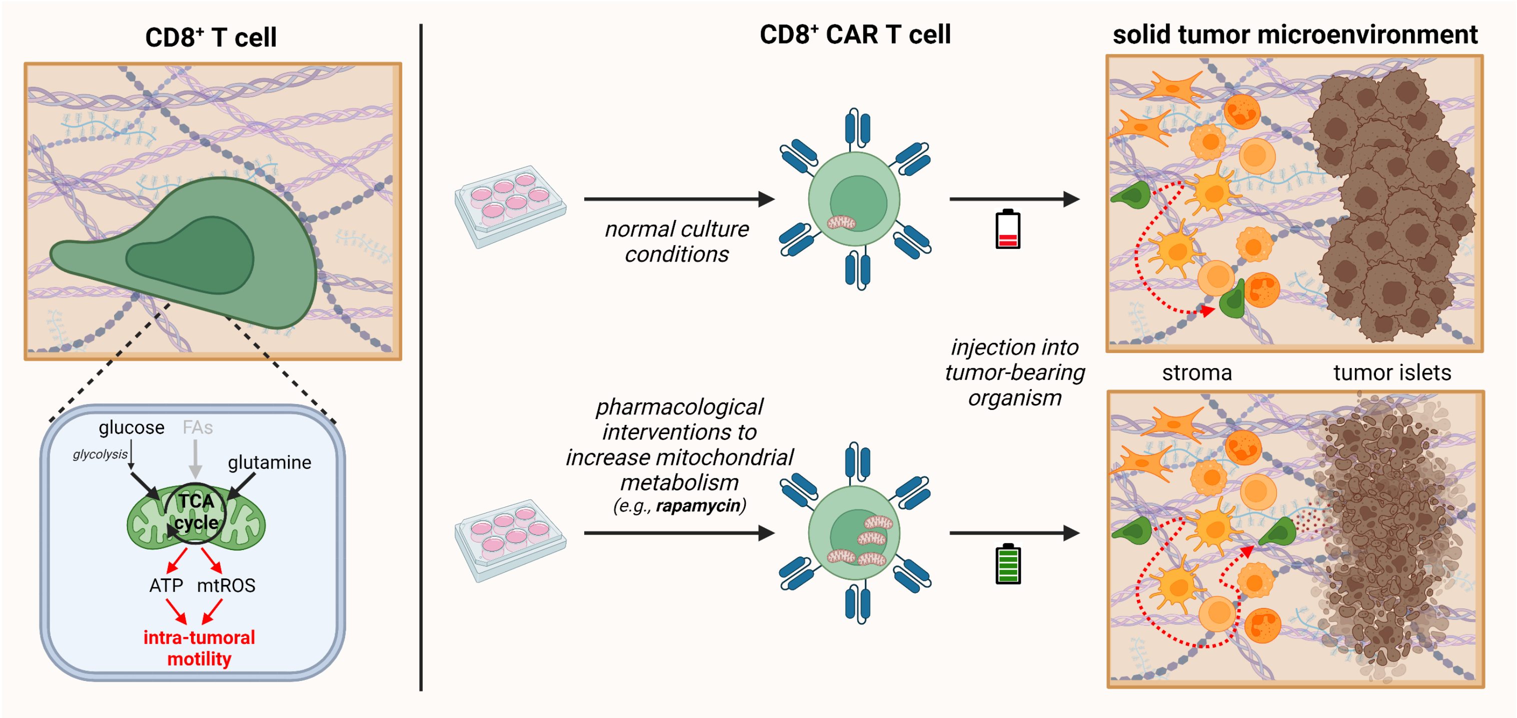


**Supplementary Figure 12. Summry of the main findings reported in the manuscript.**
On the left, TCA cycle fueled by glucose and glutamine, but not fatty acids (FAs), is the main metabolic pathways supporting human CD8$^+$ T cell migration in 3D environments, including solid tumors. Mechanistically, TCA cycle is required to support both ATP and mtROS production via OXPHOS machinery. On the right, pharmacological strategies used during *in vitro* culture to increase mitochondrial metabolism of CD8$^+$ CAR T cells are effective to confer on these cells a superior ability to infiltrate into solid tumors and reach tumor islets for efficient killing of cancer cells.
